# Allometric tissue-scale forces activate mechanoresponsive immune cells to drive pathological foreign body response to biomedical implants

**DOI:** 10.1101/2022.01.14.476395

**Authors:** Jagannath Padmanabhan, Kellen Chen, Dharshan Sivaraj, Britta A. Kuehlmann, Clark A. Bonham, Teruyuki Dohi, Dominic Henn, Zachary A. Stern-Buchbinder, Peter A. Than, Hadi S. Hosseini, Janos A. Barrera, Hudson C. Kussie, Noah J. Magbual, Mimi R. Borrelli, Artem A. Trotsyuk, Sun Hyung Kwon, James C.Y. Dunn, Zeshaan N. Maan, Michael Januszyk, Lukas Prantl, Geoffrey C. Gurtner

## Abstract

For decades, it has been assumed that the foreign body response (FBR) to biomedical implants is primarily a reaction to the chemical and mechanical properties of the implant. Here, we show for the first time that a third independent variable, allometric tissue-scale forces (which increase exponentially with body size), can drive the biology of FBR in humans. We first demonstrate that pathological FBR in humans is mediated by immune cell-specific *Rac2* mechanotransduction signaling, independent of implant chemistry or mechanical properties. We then show that mice, which are typically poor models of human FBR, can be made to induce a strikingly human-like pathological FBR by altering these extrinsic tissue forces. Altering these extrinsic tissue forces alone activates *Rac2* signaling in a unique subpopulation of immune cells and results in a human-like pathological FBR at the molecular, cellular, and local tissue levels. Finally, we demonstrate that blocking *Rac2* signaling negates the effect of increased tissue forces, dramatically reducing FBR. These findings highlight a previously unsuspected mechanism for pathological FBR and may have profound implications for the design and safety of all implantable devices in humans.

**One-Sentence Summary:** Allometric tissue-scale forces at the implant-tissue interface drive pathological foreign body response.

## Introduction

Biomedical implants have revolutionized modern medicine by improving the survival and quality of life for millions of patients worldwide. Over 70 million devices, including breast implants, pacemakers, and orthopedic prostheses, are implanted globally each year and are associated with more than $100 billion in annual expenditure (*1*). Chronic inflammation around implanted devices leads to reduced biocompatibility and results in the development of a long-term foreign body response (FBR). In clinical practice, the longevity of biomedical implants is limited by pathological FBR, frequently leading to implant failure and eventual rejection (*2*). Nearly 90% of all implant failures in commonly used medical devices are associated with FBR, and up to 30% of all implantable devices will undergo failure during their lifetime (*3–5*). As advances in materials science and electronics continue to shape the design of increasingly sophisticated biomedical devices, modifying the underlying host inflammatory response to these biomaterials remains the final frontier in developing truly biointegrative medical devices.

FBR begins as a wound healing-like response to the local tissue trauma that occurs during the initial surgical implantation of a foreign device. Shortly thereafter, FBR begins a transition toward a long-term response state in which a fibrous capsule forms around the implant, leading to both device malfunction and distortion of surrounding tissue (*2, 6*). The current prevailing hypothesis is that FBR is primarily a reaction of the local host tissue to the chemical and mechanical surface properties of the implanted material (*7–9*). Accordingly, recent research has focused on novel chemistries to identify rare, “superbiocompatible” materials such as zwitterionic hydrogels and triazole-containing alginates that appear to significantly reduce the FBR (*7, 10, 11*). Similarly, novel strategies for modulating the mechanical properties of biomaterials have also been developed, which show that soft materials can reduce fibrosis, although they do not reduce inflammation (*9*). While these developments have significantly improved our understanding of FBR, there are limitations to these approaches. Hydrogels and other soft materials have a low range of elastic moduli (1-100 kPa) and cannot be used for biomedical devices that need to provide structural support (e.g., bone and orthopedic implants) or devices that interact with relatively stiffer tissues (e.g., pacemakers and neurostimulators). Thus, the vast majority of commonly used biomedical devices continue to be fabricated from traditional materials, such as silicone and titanium, and are therefore subject to high rates of FBR-related implant failure.

Our incomplete understanding of FBR is exacerbated by the inability of standard laboratory models to recapitulate the sustained inflammatory response and severe fibrotic reaction associated with implant failure in humans (*12*). Even though the molecular machinery responsible for inflammation and fibrosis is highly conserved across species, studies have shown that small animal models, such as mice, generate a relatively mild FBR to implantable materials compared to humans (*13, 14*), which limits their clinical relevance. A key feature that differentiates humans from mice is their body size, with humans being several orders of magnitude larger (*15, 16*). Well-established allometric scaling principles dictate that tissue-scale forces and, thus, tissue mechanical stress increase exponentially with an increase in body size (*15–18*). While it seems likely that these changes in allometric tissue-scale forces could play a role in inter-species differences in FBR, these mechanisms have not been examined before.

Here, we comprehensively investigate FBR in both human and murine models and determine how changes in allometric tissue forces may affect the development of FBR. Using this knowledge, we then manipulate extrinsic tissue forces using a novel murine model to recapitulate all aspects of human FBR on both a histological and transcriptomic level. In both humans and mice, we identify a unique subpopulation of mechanoresponsive myeloid cells mediated by RAC2 signaling that specifically respond to changes in tissue forces during FBR. Further, we demonstrate that pharmacological inhibition of these cell populations can therapeutically prevent the development of pathological FBR.

## FBR in humans is characterized by similar fibrotic encapsulation across a diverse array of implants, regardless of implant chemistry or mechanical properties

To understand the importance of material properties on human FBR, we analyzed fibrotic capsules from a diverse array of biomedical implants. We collected human fibrous capsule tissue samples from silicone-based breast implants, titanium-based pacemakers, neurostimulators, and mixed alloy-based orthopedic implants. To our knowledge, we provide the first reported comparison of human FBR across diverse implant types using histological analyses. Since each type of implant was made up of different material chemistries with different mechanical properties, we hypothesized that the resultant fibrotic capsule surrounding each type of implant would vary in fibrotic severity.

Surprisingly, we found that the FBR surrounding each implant was strikingly similar in tissue architecture (**Fig. 1A-B; Fig. S1**). Specifically, all FBR capsules analyzed were predominantly composed of mature type I collagen with highly organized and aligned fibers, characteristic of severely fibrotic scar tissue resulting from elevated tissue mechanical forces (**Fig. 1C,D; Fig. S1**) (*19*). Overall, these implants all had different material chemistries with different mechanical properties, yet they all generated similar levels of FBR development. Thus, we postulated that implant material properties were not sufficient to explain the mechanisms that underlie human FBR.

**Figure 1.**
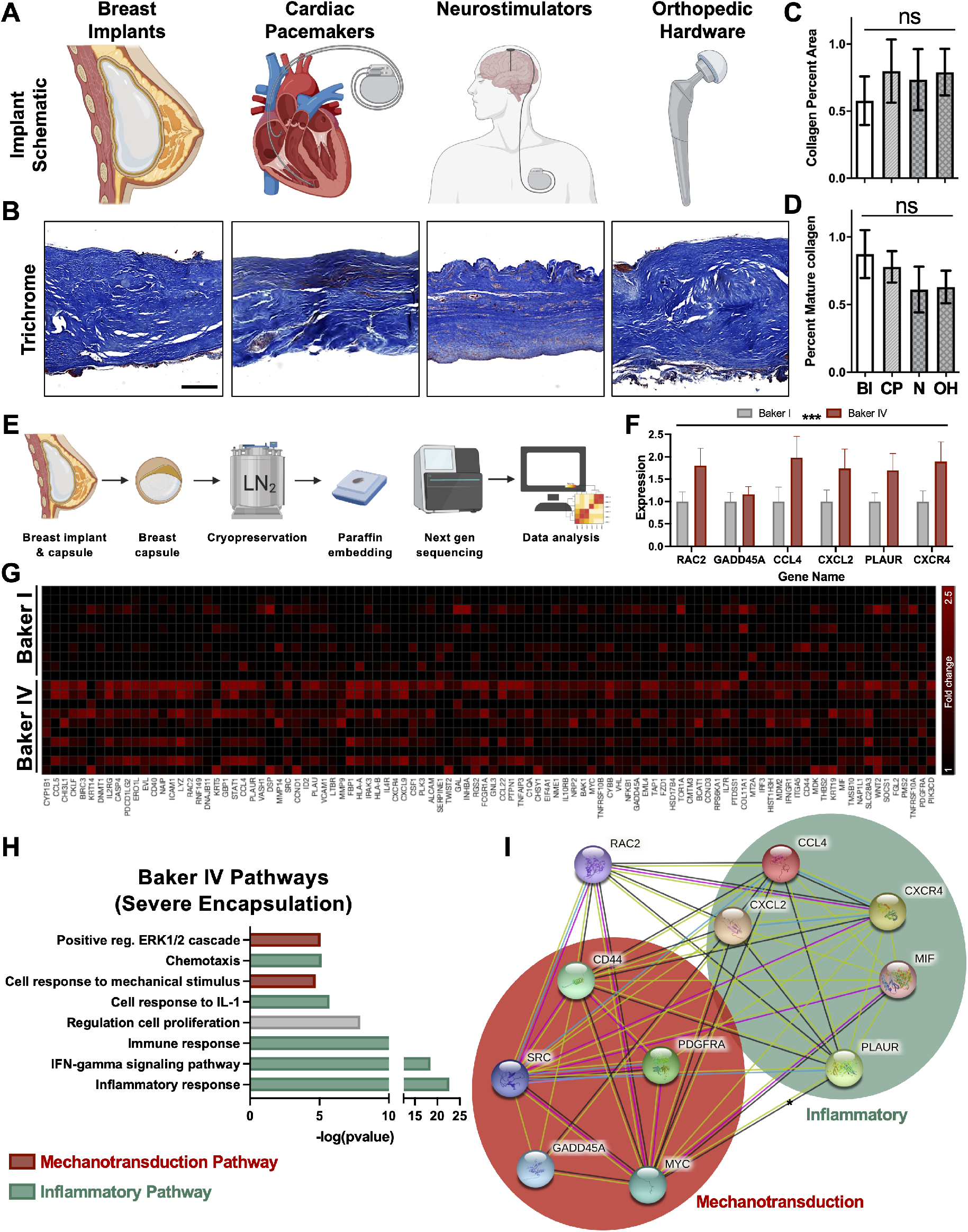
Pathological FBR in humans is mediated by *RAC2* mechanotransduction signaling, regardless of implant properties and is associated with increased mechanical signaling. **(A)** Schematic of various implant types. **(B)** Trichrome staining of fibrotic capsules from the fibrous capsule formed around silicone-based breast implants, titanium-based pacemakers and stainless steel-based orthopedic implants are all strikingly similar to one another. (n=4-6) for each implant category. **(C,D)** Quantification of collagen and mature collagen shows no significant differences between the different types of human implants. BI = Breast Implants; CP = Cardiac Pacemakers; N = Neurostimulators; OH = Orthopedic Hardware. **(E)** Schematic showing the experimental methodology followed; FBR capsules from Baker I and Baker IV breast implants were subject to molecular analyses using a commercially available biomarker panel (HTG Molecular). A total of 9 Baker I specimen and 11 Baker IV specimen were used in this study. **(F)** Selected mechanotransduction and inflammatory genes upregulated in Baker IV (red) versus Baker I (grey) capsules. **(G)** Heatmap of the top 100 genes upregulated in Baker IV vs. Baker I breast implants, organized in decreasing order of fold change from left to right. **(H)** Pathways significantly upregulated in Baker IV samples analyzed using Database for Annotation, Visualization and Integrated Discovery (DAVID). Pathways highlighted in red are mechanical signaling pathways and those highlighted in green are inflammatory pathways. **(I)** STRING (Search Tool for the Retrieval of Interacting Genes/Proteins) analysis showing that *Rac2* is a central mediator of both mechanotransduction signaling as well as inflammatory signaling genes that were all upregulated in Baker IV specimen. STRING analyzed interactions between the different genes based on experimental evidence as well as the predicted interactions based on various databases, which are color-coded as listed below: pink (experimentally determined), blue (curated databases), green (gene neighbourhood), red (gene fusion), blue (gene co-currence), light green (textmining), black (co-expression), violet (protein homology).

## Pathological FBR in humans is characterized by RAC2 mediated mechanotransduction and inflammatory signaling

To explore other, previously unidentified variables that may be involved in FBR, we analyzed implant capsules from identical biomedical breast implants that still generated different severities of foreign body response (FBR) in human patients (*20*). Fibrosis around breast implants is conventionally classified using the Baker system, where Baker I is the least severe and represents cases with minimal clinically observable implant capsule contracture, while Baker IV represents the most severe (pathological) cases that display a sustained inflammatory reaction, pronounced fibrotic contracture, and pain. Since relatively few patients with mild (Baker I) capsular fibrosis routinely undergo surgical revision and tissue collection, this group represented a limiting factor in our overall data collection. We analyzed mRNA isolated from human tissue specimens of both mild (non-pathological) and severe (pathological) FBR observed in humans around implants made of the exact same silicone material using a next generation sequencing-based quantitative assay against a biomarker Panel (HTG Molecular), which consisted of 2500+ known biomarkers for inflammation and fibrosis (**Fig. 1E-G**).

We displayed the top genes that were significantly (*p<0.05) upregulated in the Baker IV breast implant capsules compared to Baker I capsules in a heatmap (**Fig. 1G**) and then used the Database for Annotation, Visualization, and Integrated Discovery (DAVID) pathway analysis to identify the top pathways upregulated in the severe Baker IV compared to Baker I capsules (**Fig. 1H**) (*21*). We observed that the top genes upregulated in Baker IV implants were critically involved in mechanical signaling pathways including “*cellular response to mechanical stimulus*” and “*positive regulation of Erk1 and Erk2 cascade*” (**Fig. 1H**). Baker IV implant capsules also showed significant upregulation of “*inflammatory signaling related to chemotaxis*”, “*cellular response to interleukin-1*”, and “*immune response pathways*” (**Fig. 1H**). In contrast, the milder Baker I implant capsules displayed only a modest upregulation of pathways related to homeostatic processes such as glucose and fat metabolism (**Fig. S2**).

Since prior research has shown that elevated mechanical force subsequently activates inflammatory signaling (*22*), we postulated that upregulation of upstream mechanotransduction signaling genes might drive cellular responses towards Baker IV inflammation, fibrosis, and pathologically severe FBR. Based off this, we found several critically upregulated mechanotransduction and inflammatory genes within our human dataset (**Fig. 1F**). Most interestingly, we found that *RAC2*, a hematopoietic-specific Rho-GTPase inflammatory mechanotransduction marker, was significantly upregulated in the Baker IV implant specimens (**Fig. 1F, G**). *RAC2* is a signal transduction molecule, which mediates the recruitment and activation of immune cells and has been shown to be activated by mechanical forces (*23, 24*). Baker IV capsules also demonstrated increased expression of *GADD45A,* a mechanoresponsive gene that mediates inflammation and tissue response to injury (*25, 26*), and *CCL4*, which has been shown to be mechanically activated and mediates classical fibrosis through the activation of hematopoietic cells (*27–29*). With respect to inflammatory pathways, Baker IV capsules significantly upregulated *CXCL2, PLAUR, MIF*, and *CXCR4*. Of these, *CXCL2* and *CXCR4* have been shown to be directly activated by *RAC2*, contributing to the recruitment and activation of myeloid cells including neutrophils and macrophages (*30–32*). Similarly, *MIF* and *PLAUR* have also been identified as critical mediators of inflammation in hematopoietic cells, leading to fibrosis (*33, 34*).

Using STRING (Search Tool for the Retrieval of Interacting Genes/Proteins), a pathway analysis tool that predicts gene-gene interactions (*35*), we found that *RAC2* played a centralized and pivotal role, guiding the expression of all other aforementioned top genes to drive both mechanotransduction and inflammatory pathways (**Fig. 1I**). Additionally, *RAC2* guided the activation of *CD44*, which has been previously shown to increase the responsiveness of the extracellular matrix to mechanical stimulation and described to be crucial for the activation of downstream effectors such as *SRC* and *MYC*, both of which directly contribute to macrophage-mediated inflammation (*36–38*).

Since both Baker I and Baker IV implants are made of the same material (silicone) with identical material chemistry and mechanical properties, these results strongly suggested that pathological (Baker IV) FBR in humans may be mediated by *RAC2* mechanotransduction signaling, independent of implant material properties. Since RAC2 is a hemopoietic-specific Rho-GTPase (*23, 24*), we hypothesized that mechanical forces may mediate immune cell-specific mechanotransduction to generate pathologically severe FBR. We then sought to test this hypothesis in a novel murine model.

## Standard murine models cannot recreate the high mechanical stress environment surrounding human implants due to allometric tissue scaling properties

To further study the importance of mechanical forces on the development of FBR, we decided to utilize a murine model of FBR. Unfortunately, previous studies have shown that mice generate a relatively mild FBR to implantable materials compared to humans (*13, 14*), leading to a dearth in the development of clinically relevant therapies for FBR. Because of the dramatic difference in mechanical signaling observed between severe (pathological) and mild (non-pathological) FBR in humans, we hypothesized that increased mechanical signaling due to allometric tissue-scale forces might also explain differences in FBR severity between mice and humans. In the context of biomedical implants, cells are believed to mediate mechanotransduction signaling primarily due to the implant material properties. For example, cells on stiffer substrates have been found to exhibit higher mechanical signaling compared to cells cultured on softer substrates (*39, 40*). However, as organisms evolve and grow larger, fundamental allometric scaling principles dictate that mice have inherently exponentially lower levels of extrinsic tissue forces compared to humans owing to the 10^4^-fold difference in organism size (*15–18*). These drastic differences in the surrounding host tissue properties would significantly affect the mechanical environment surrounding the implant. Thus, we must consider both the implant materials properties and surrounding tissue properties to quantify the local mechanical stress at the implant-tissue interface.

To investigate how differences in murine and human extrinsic tissue-scale forces play a role in the development of FBR, we modeled the local mechanical stress patterns at the implant-tissue interface in mice and humans using finite element modeling (FEM) (ABAQUS, version 2017, SIMULIA, Providence, RI). Utilizing a combination of factors, including the murine or human surrounding tissue properties and the material properties of the implants themselves (**Methods; Fig. S3**), we predicted that the maximal stress surrounding a standard silicone murine implant was 0.2 kPa, compared to over 20 kPa surrounding human silicone breast implants (**Fig. 2A; Fig. S4**). Since both implants were made of the same material (silicone), this 100-fold increase in mechanical stress in humans was largely due to the allometric differences between humans and mice in both tissue size and tissue mechanical properties, potentially explaining why standard murine preclinical models generate much lower FBR (**Fig. 2A; Fig. S3, S4**). Correspondingly, we found that humans generated FBR with significantly increased amounts of collagen and mature collagen compared to standard murine models (*p<0.05; **Fig. 2C-G**). Human implant capsules were also characterized by significantly increased myofibroblast activation compared to standard murine implants (*p<0.05; **Fig. 2E**). Analyzing the surface of implants using scanning electron microscopy (SEM), we also observed that human implants were covered by a highly fibrous collagen network, which was not observed on the standard murine implants (*p<0.05; **Fig. 2F, G**).

**Figure 2.**
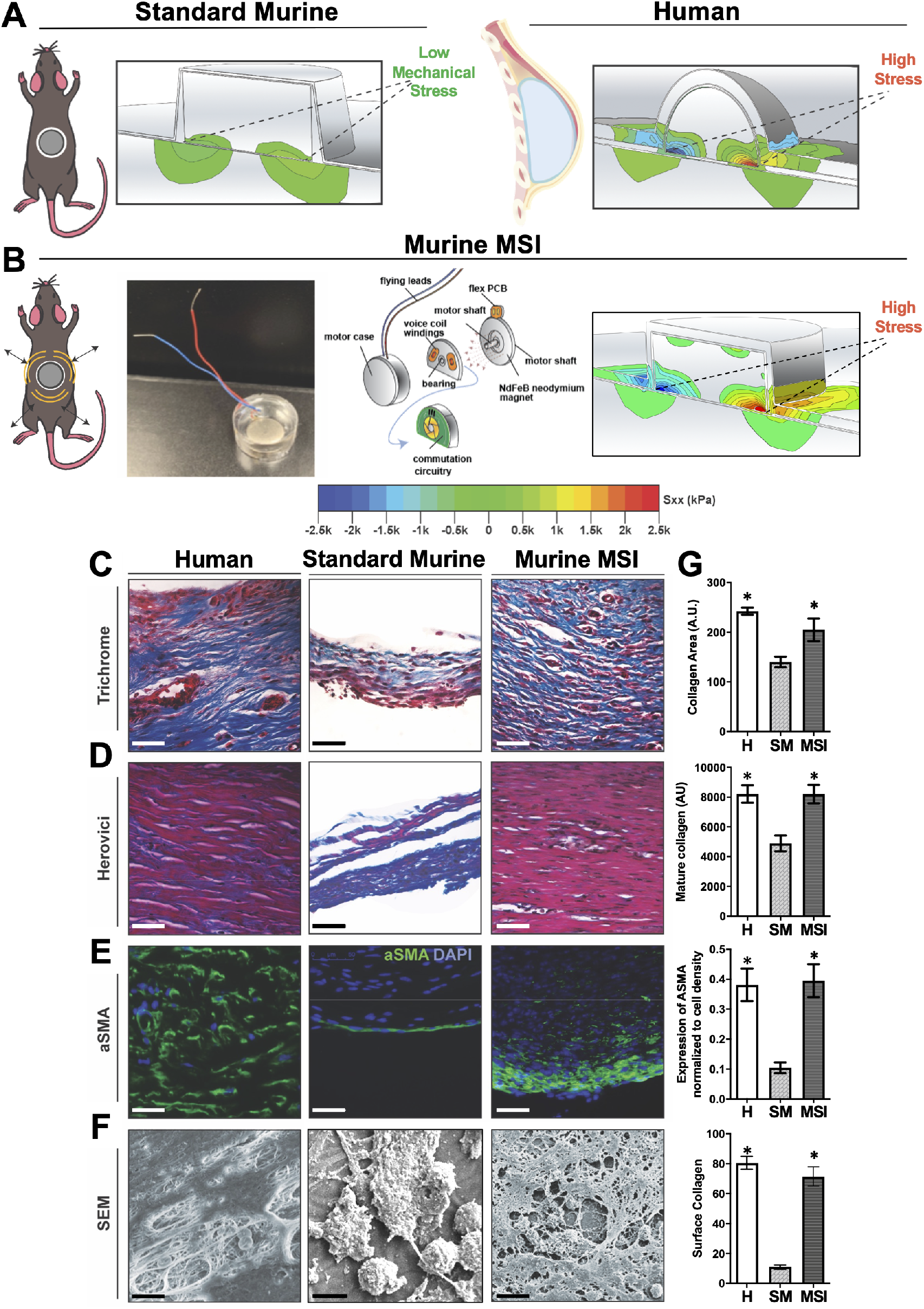
Altering extrinsic tissue-scale forces using mechanically stimulating implants (MSIs) produces human-like FBR capsule architecture in mice. **(A)** Finite element modeling of murine and human implants showing that human implants are subject to 100-fold higher mechanical stress than murine implants. **(B)** Schematic and picture of the MSI model. FE modeling confirming that MSI recreate human-levels of mechanical stress in the mouse. **(C)** Trichrome staining of FBR capsules in the human implant capsules, standard murine model, and the MSI murine model. Scale Bar = 50 μm. **(D)** Herovici staining showing mature (red) and immature (blue) collagen. Scale bar = 50 μm. **(E)** Immunostaining for alpha smooth muscle actin (αSMA), a marker for myofibroblasts. Scale bar = 50 μm. **(F)** Scanning electron microscopy (SEM) imaging of the surface of the capsules. Scale bar = 10 μm. **(G)** *Top row*: Quantification of percent area positive for collagen in each capsule (far right column). n=5 for each group. *p < 0.05. *Second row*: Quantification of mature collagen deposition in the FBR tissue (far right). n=5 for each group. *p < 0.05. *Third row*: Quantification of αSMA normalized to cell density using image analyses in each capsule. *Fourth row*: Quantification of surface collagen percent area associated with each capsule. n=8 images for each group. *p < 0.05. H = Human; SM = Standard Murine.

Interestingly, variations in implant geometry and implant material stiffness resulted in only minimal changes to the predicted mechanical stress at the implant-tissue interface in our finite element models. In humans, both silicone-based breast implants and much stiffer titanium-based implants, such as pacemakers and neurostimulators, generated similar mechanical stress profiles in the range of 20-23 kPa surrounding the implants (**Fig. S4**). These similarities in stress profiles could explain the similar fibrotic encapsulation observed around these different types of human implants (**Fig. 1A-D**). Overall, by controlling for all other factors including implant material properties, we found that the large differences in murine and human extrinsic tissue mechanical forces significantly contributed to drastically different mechanical stress profiles surrounding biomedical implants.

## Increasing extrinsic tissue-scale forces using mechanically stimulating implants (MSIs) produces human-like FBR capsule architecture in mice

To investigate the importance of these tissue forces, we developed a novel murine model to artificially impose human-like levels of increased tissue mechanical forces surrounding implants, independent of implant size or chemistry. Specifically, we developed an implantable silicone device encapsulating a small motor (**Fig. 2B; Fig. S5, S6**), which could be induced to produce intermittent *in situ* implant vibration. Since vibration is a mechanical force, these mechanical forces from the implant would intermittently increase the mechanical loading of the surrounding tissue to human levels. After iterating through combinations of vibration frequency and amplitude in our finite element model, we determined that 3V batteries with an amplitude of 1.38G and a frequency of 203 Hz would artificially increase the extrinsic tissue-scale forces from the surrounding host tissue to generate a 100-fold increase in mechanical stress at the implant-tissue interface (24.1 kPa), similar to that surrounding human implants (**Fig. 2B**). The development of these mechanically stimulating implants (MSIs) in mice provided a unique platform to examine the effect of increased extrinsic tissue-scale forces alone on the subsequent foreign body response.

When compared to standard implants, MSIs developed FBR capsules with significantly more collagen deposition in the capsule and on the implant surface, higher collagen maturity, and upregulated myofibroblast activation (*p<0.05; **Fig. 2C-G**). Accordingly, MSIs developed FBR capsules that were nearly identical to human FBR capsules across all these metrics (**Fig. 2C-G**). These findings demonstrated that by artificially inducing high levels of mechanical stress around murine implants to imitate the mechanical environment in humans, a human-like FBR capsule architecture can be recreated in mice, which is markedly different than that observed in standard murine implants (*12, 41*).

Both the MSI and standard implants were made of the same material (silicone) and had the same geometry. To control for any differences resulting from the presence of the inactivated coin motor, we performed additional experiments to compare the FBR between MSIs with the motor on and off (**Fig. S7**). We found that the MSIs without the motor activated were also unable to generate a human-like highly fibrotic capsule (**Fig. S7**), confirming that the human-like FBR observed with the motor activated MSIs (producing vibration) was entirely due to mechanical loading of the surrounding tissue. Thus, we show that increased extrinsic mechanical forces by the surrounding tissue results in a remarkably human-like highly fibrotic capsule, independent of both material chemistry and mechanical properties (**Fig. 2; Fig. S7**). While elevated mechanical tissue forces have been previously linked to increased inflammation and fibrosis in the context of wound healing, we demonstrate this phenomenon for the first time in the context of FBR to biomedical implants (*42*). Taken together, these results show for the first time that manipulating a third independent variable, the extrinsic tissue-scale forces (which can increase either allometrically in humans or artificially with vibration in mice), can drive the biology of FBR, independent of implant properties.

## Elevated extrinsic tissue-scale forces promote sustained activation of immune cell-specific *Rac2* mechanotransduction to drive human-like pathological FBR in mice

To examine the cellular response through which extrinsic tissue-scale forces alter FBR, we analyzed the cells surrounding the murine MSI and standard implants using single cell RNA sequencing (scRNA-seq). We analyzed a total of 36,827 cells from both early-stage and late-stage capsules from both standard implants and MSIs (**Fig. 3A**). These time points were chosen due to observable differences in the tissue architecture of the capsules as early as 2-weeks post-implantation (early-stage), with the MSI capsules progressing to reach a stable human-like tissue architecture at about 4-weeks post-implantation (late-stage) (**Fig. 2; Fig. S7**).

**Figure 3.**
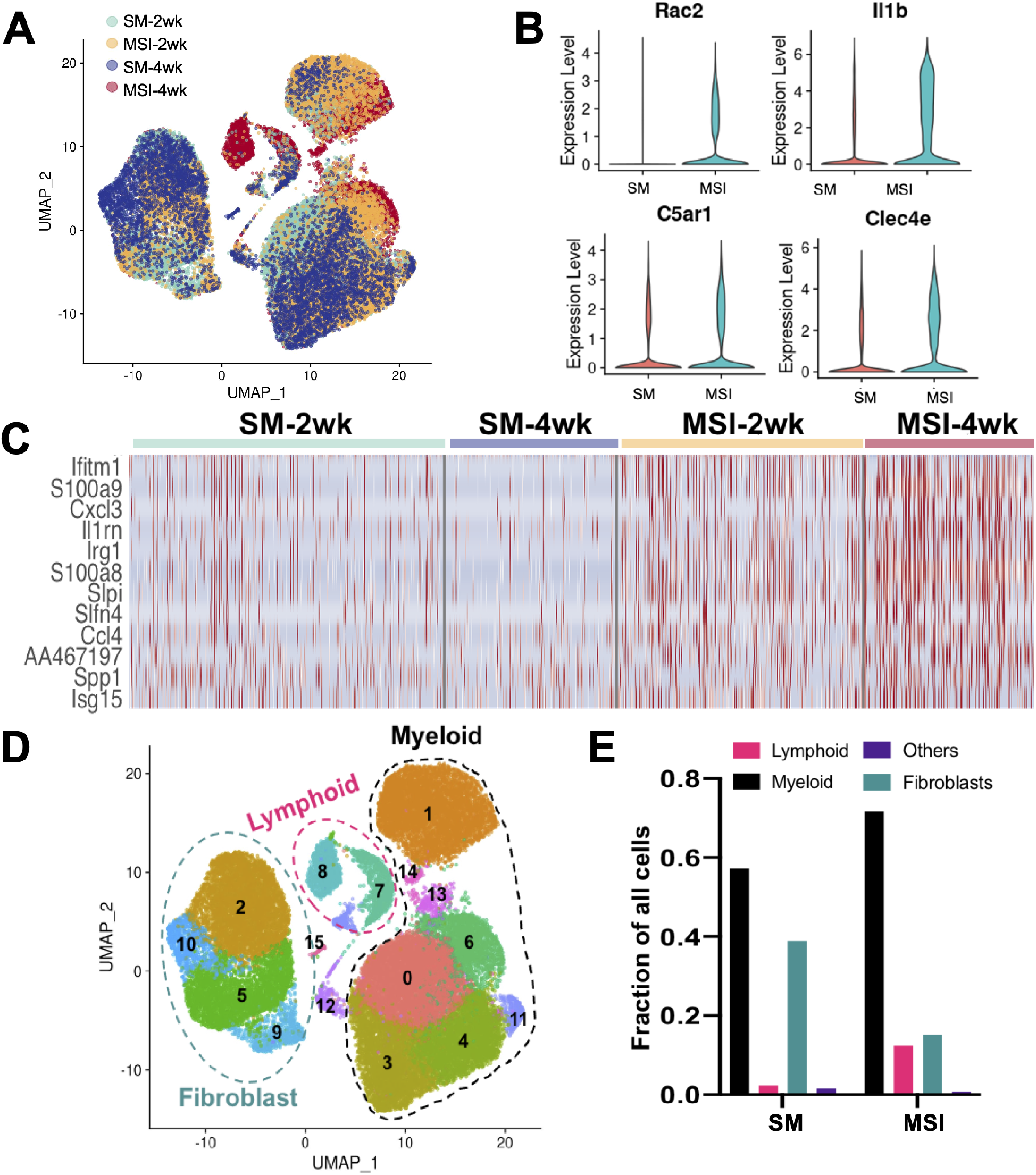
MSIs generate a sustained inflammatory response at the implant-tissue interface. **(A)** UMAP plots of all cells from murine FBR capsules classified by sample type and timepoints. A total of 36,827 cells were analyzed. **(B)** MSI capsules show a robust activation of *Rac2* and associated inflammatory markers as compared to standard implant capsules. **(C)** Heatmap of differentially regulated inflammatory genes between standard implant cells and MSI cells, showing a similar pattern of expression. All inflammatory genes analyzed are upregulated in MSI capsules at both early and late timepoints. In contrast, most inflammatory markers show a modest activation in standard implants at the early timepoint, which subsides at the late timepoint. **(D)** UMAP plots of all cells from murine FBR capsules classified by Seurat clusters. Two major cell types were found in both standard implants and MSIs: immune cells (myeloid & lymphoid) and fibroblasts. **(E)** Relative proportion of myeloid, lymphoid and fibroblast cells in standard murine implants and MSI capsules. Myeloid cells were the most abundant cell type in both capsules and were especially enriched with mechanical stimulation.

Analysis of the differential gene expression between cells isolated from standard implant capsules and MSI capsules revealed significant differences in the activation of *Rac2* and associated inflammatory markers between the two groups. MSI cells showed robust activation of *Rac2* signaling in contrast to standard implant cells, which showed relatively minor activation of *Rac2* (**Fig. 3B**). Similarly, MSI cells showed robust activation of inflammatory markers such as *Il1b, Clec4b, and C5ar1* (**Fig. 3B**) (*43–45*). Standard implants initially expressed a modest activation of these inflammatory markers, which subsided at later timepoints, while MSI capsules showed a robust early activation of these markers, which continued to increase in later time points (**Fig. S8A**). This pattern of expression could be seen in a heatmap of expression of top inflammatory genes, such as *Cxcl3* and *Ccl4*, in which cells increased expression in MSI models and over time (**Fig. 3C; Fig. S8B**). Thus, our murine MSI model recapitulated the overall upregulation of *Rac2* signaling and inflammation induced by mechanical stimulus that was observed in the transcriptomic profiles of human Baker IV severely fibrotic capsule tissue (**Fig. 1**).

After establishing that our MSI model recapitulates the Rac2 mediated mechanotransduction environment observed in humans, we then proceeded to characterize the critical cell types that drive FBR. Automated cell identification with SingleR revealed two major cell types that populated FBR capsules in both standard implants and MSIs: immune cells (myeloid and lymphoid) and fibroblasts (**Fig. 3D, E**) (*46*). Myeloid cells, including monocytes, macrophages, dendritic cells, and granulocytes, were the most abundant cell type in both the standard implant and MSI capsules (**Fig. 3D, E**), and mechanical stimulus directly increased myeloid cell proportions in the MSI model (**Fig. 3E**).

## *Rac2* immune signaling drives Baker IV FBR

We separately analyzed each major cell type (i.e., myeloid, lymphoid, and fibroblasts) to determine cell-type specific transcriptional shifts induced by increased levels of extrinsic tissue-scale forces. We first focused our analysis on myeloid cells, since they made up the majority of the capsules and seemed to directly increase with mechanical stimulus (**Fig. 3D, E**). Myeloid cells clustered in 8 distinct subpopulations (Clusters 0-7; **Fig. 4A**). Of these myeloid cell clusters, Clusters 1 (granulocytes) and Clusters 4&7 (macrophages) were specifically enriched in MSIs (**Fig. 4B**).

**Figure 4.**
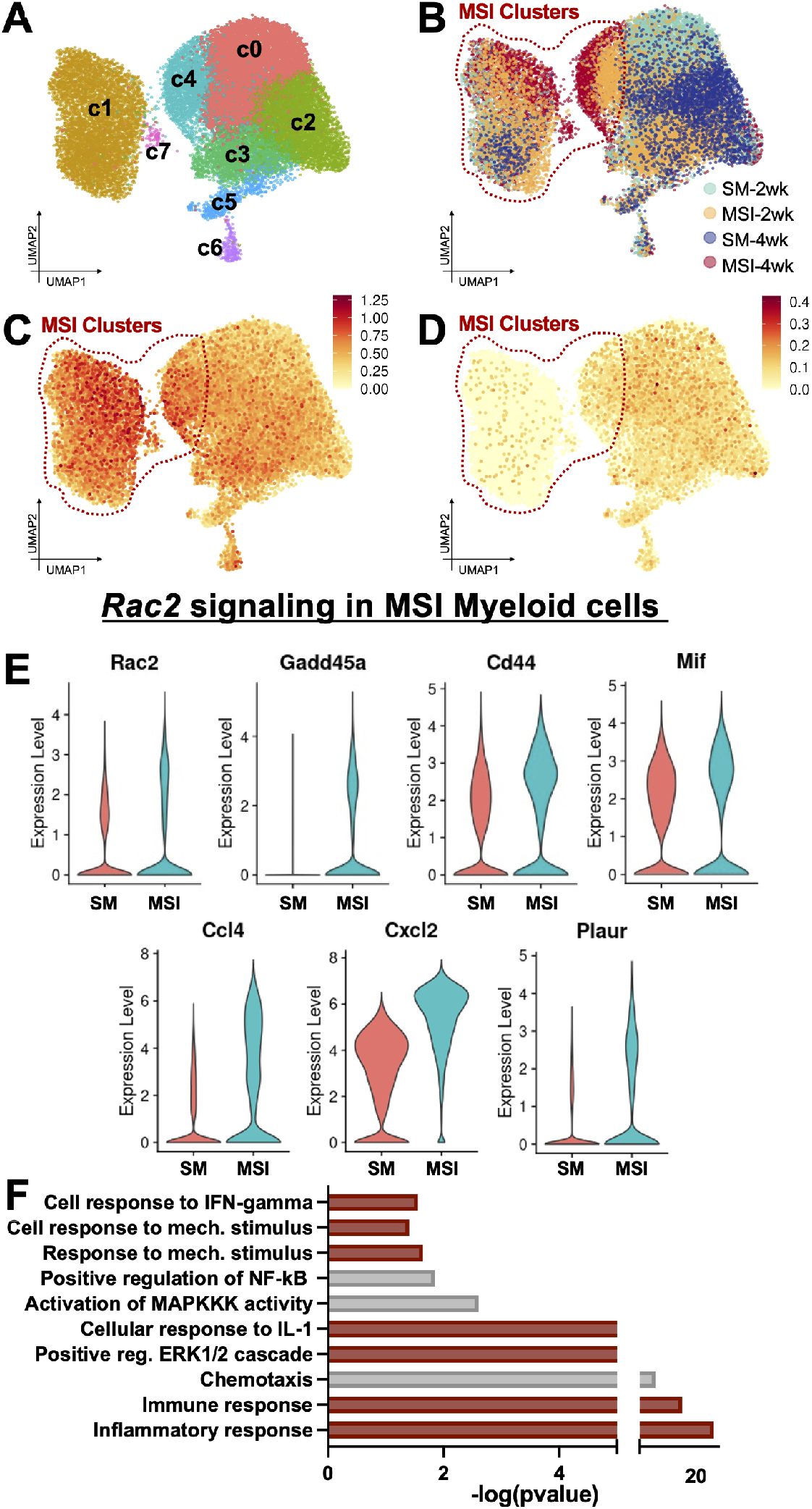
Increased extrinsic tissue-scale forces activate *Rac2* signaling in myeloid cells, which drives a Baker IV fibrotic phenotype in mice. **(A,B)** UMAP Plot of myeloid cells in SM and MSI implant capsules. Clusters 1,4,7 (red dotted circle) are highly enriched in MSI capsules. **(C,D)** FeaturePlot of top averaged **(C)** Baker IV markers (Fig. 1B), including key mechanotransduction and inflammatory chemokine signaling pathways, and **(D)** Baker I markers. **(E)** Violin plots of Baker IV markers differentially upregulated in the MSI clusters. **(F)** Pathways significantly upregulated in Baker IV samples analyzed using Database for Annotation, Visualization and Integrated Discovery (DAVID). Pathways highlighted in red are also upregulated in Baker IV human specimen.

Next, we examined how the transcriptional profiles of these murine clusters compared to the expression of Baker IV human biomarkers (**Fig. 1**), i.e., the genes that were highly upregulated in Baker-IV (pathological) human specimens. We determined the combined mean expression of the top 25 Baker IV biomarkers (**Fig. 1G**), which included mechanical signaling and downstream inflammatory genes, to create a “gene signature” that represented the transcriptomic profile of pathological, severe human FBR. Plotting the expression of this gene signature (**Fig. 4C**), we found that genes related to Baker IV pathological FBR were highly upregulated in MSI clusters (Clusters 1,4,7) compared to expression in standard implant clusters (**Fig. 4B, C**). We then created a gene signature of the top 25 Baker I biomarkers and found that standard implant clusters instead upregulated genes found in the mild human Baker I capsules (**Fig. 4D**), which indicated normal homeostatic processes (*47, 48*).

In addition, *Rac2* and downstream mechanotransduction genes (e.g., *Gadd45a, Mif, Cd44)* and inflammatory genes (e.g., *Cxcl2, Ccl4, Plaur*) were also differentially upregulated in the MSI clusters, as with the top differential genes of interest observed in human Baker IV capsules (**Fig. 1E,F; Fig. 4E**) (*33, 49, 50*). DAVID pathway analyses further confirmed the significant overlap in the gene expression between MSI cells and human Baker IV capsules (**Fig. 4F**). Both human baker IV and murine MSI cells upregulated key mechanotransduction pathways, including “*Positive regulation of Erk1 and Erk2 cascade*” and “*Cellular response to mechanical stimulus*”, as well as inflammatory pathways, such as “*Interferon-gamma signaling*” and “*Chemotaxis*” (**Fig. 1H; Fig. 4F**).

Overall, these myeloid cells expressed a wide array of genes and pathways with significant overlap with human FBR transcription, with mechanical stimulation of immune cells promoting a pathological Baker IV profile and standard murine implants promoting a benign Baker I profile. The differential upregulation of inflammatory markers along with the increased presence of myeloid cells in MSI capsules further demonstrated that the human-like FBR in MSIs is characterized by a mechanically-induced and sustained inflammatory response at the implant-tissue interface.

## Elevated tissue-scale forces reproduce all classic features of pathological FBR by activating fusogenic macrophages, MHC Class II lymphocytes, and myofibroblasts

Fusion of macrophages leading to the formation of foreign body giant cells (FBGCs) is a hallmark of the classic FBR (*6*). Fusogenic macrophages and FBGCs are known to release degradative enzymes, ROS, and pro-fibrotic factors, which regulate the recruitment, growth, and proliferation of fibroblasts. We found that *Arg1^+^* macrophages (Cluster 4), which have been previously reported to be fusogenic macrophages, were highly enriched in MSI capsules as compared to standard implants (**Fig. 5A**) (*51*). It was recently reported that *Rac1*-mediated fusion of macrophages requires the expression of *Mmp14* and that *Mmp14^null^* cells do not fuse (*52*). Cluster 4 macrophages showed a simultaneous upregulation of *Arg1, Rac1*, and *Mmp14* (**Fig. 5A**), demonstrating the activation of fusogenic macrophages in MSIs in response to increased tissue-scale forces at the implant-tissue interface.

**Figure 5.**
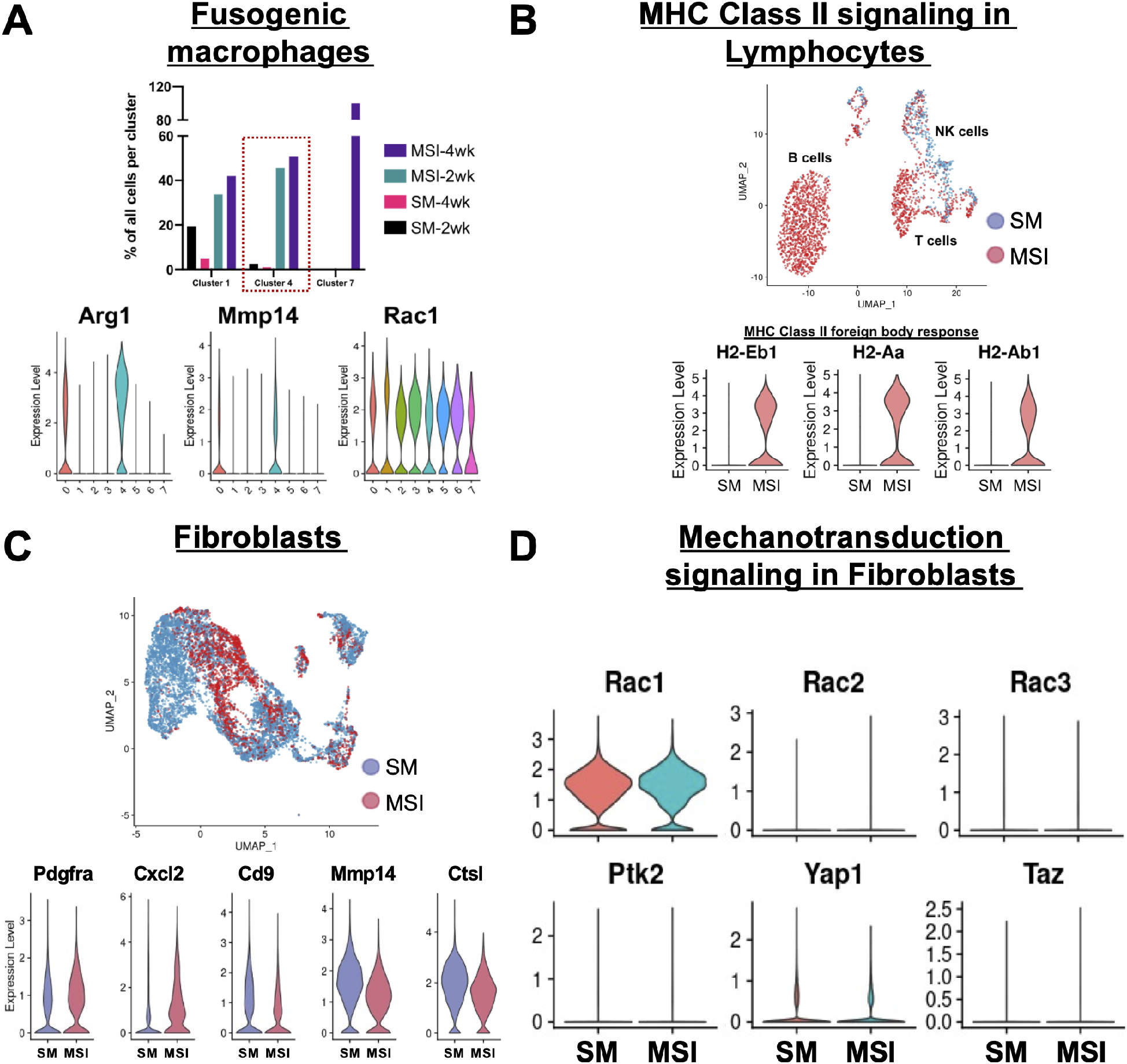
Increasing tissue-scale forces activates fusogenic macrophages, MHC II lymphocytes and myofibroblasts, all classic features of pathologic FBR. **(A)** Relative proportions of cell types in Clusters 1,4,7 cells. Cluster 4 cells upregulate markers for fusogenic macrophages (red dotted line). **(B)** UMAP and Violin plots of lymphocytes from murine FBR capsules. MSI lymphocytes show upregulation of MHC Class II signaling. **(C)** UMAP and Violin plots of fibroblasts from murine FBR capsules. **(D)** Violin plots of mechanotransduction markers in SM and MSI fibroblasts.

Since macrophage fusion is promoted by activated lymphocytes, we next analyzed all lymphocytes in our dataset to identify the lymphoid-specific changes in transcriptional activity induced by mechanical stimulation. MSI lymphocytes showed a preferential upregulation of MHC class II foreign body response components including *H2-Eb1, H2-Aa*, and *H2-Aa1* (**Fig. 5B; Fig. S9**). Since MHC class II signaling in lymphocytes is a well-known promoter of macrophage fusion and the overall foreign body response, this provided evidence that the mechanical activation of immune cells then signaled macrophage fusion to contribute to increasing severity of FBR (*6, 53, 54*).

As expected, we also observed the presence of fibroblasts in the implant capsules. Historically, fibroblasts are thought to be critical in FBR since these cells are responsible for synthesizing collagen that crosslinks in the extracellular space and contributes to the formation of a dense, collagen-rich fibrotic capsule around the implant (*55*). We observed that several inflammatory cytokines linked to the activation of fibroblasts, such as *Cxcl2, Plaur,* and *Ccl4* (*33, 49, 50, 56, 57*), were upregulated in MSI capsules (**Fig. 4; Fig. S9**). We then compared fibroblasts from both implant models (**Fig. 5C Fig. S10**) and found that fibroblasts in the standard murine model demonstrated upregulation of proteolysis (*Mmp14, Ctsl*) and negative regulation of cell proliferation (*Cd9*), suggesting a resolving phenotype (*58–60*). In contrast, MSI fibroblasts showed an upregulation of myofibroblast marker *Pdgfra*, profibrotic cytokines such as *Cxcl2*, and downregulation of anti-proliferation marker *Cd9* (**Fig. 5C**) (*58, 61, 62*), indicative of a more activated fibroblast phenotype in MSIs. This led to the increased differentiation of myofibroblasts and collagen deposition in MSI capsules, consistent with our observations of increased myofibroblasts populating the MSI capsules (**Fig. 2E**).

Surprisingly, fibroblasts from MSI capsules and standard implant capsules showed minimal activation of canonical fibroblast mechanotransduction genes such as *Ptk2, Yap1*, and *Taz* (*17, 55*). In addition, fibroblasts displayed almost identical levels of *Rac1* activation and minimal activation of *Rac3*. Since *Rac2* is a hematopoietic-specific marker (*63–65*), fibroblasts did not express *Rac2*, further increasing our confidence that immune cells are the primary cell type responsible for the initial *Rac2* mediated mechanotransduction to drive FBR. Since *Rac2* expression is differentially upregulated in FBR capsules and myeloid cells make up the majority of cells in chronic FBR capsules, immune cells serve as a primary mechanosensor in both murine and human FBR. Thus, it appears that although fibroblasts produce collagen to create fibrotic tissue, the activation of fibroblasts is primarily mediated by activating these immune cells in the context of FBR.

## Blocking *Rac2* signaling negates the effect of increased tissue forces, dramatically reducing FBR

Our findings show that allometric tissue-scale forces activate *Rac2* signaling in immune cells, which drive the classic human pathological FBR. Since the extrinsic tissue-scale forces are inherent to the size of the organism and cannot be altered, it would require pharmacological strategies to block the mechanical activation of immune cells in FBR. To this end, we tested the efficacy of a small molecule Rac inhibitor (EHT 1864 2HCl) (*66*) in reducing FBR in our MSI model (**Fig. 6**). Local injection of EHT 1864 2HCl in the MSI model reduced the expression of immune cell-specific *Rac2* in the FBR capsules by about 80%, indicating a significant reduction in the recruitment and activation of mechanoresponsive immune cells (**Fig. 6A**). Correspondingly, we observed a significant reduction in the activation of myofibroblasts in MSI capsules treated with the small molecule inhibitor by about 90% (**Fig. 6B**). Blocking *Rac2* signaling in mice significantly reduced the overall FBR as well, specifically demonstrated by decreased capsule thickness and collagen deposition (**Fig. 6C, D**). Taken together, these results show that blocking *Rac2* signaling in immune cells can cause a cascade of downstream effects, including significantly decreased myofibroblast differentiation, reduced downstream collagen production, and mitigated FBR capsule formation. In short, by blocking the immune orchestrators of FBR, it is possible to reverse the human-like FBR resulting from increased levels of extrinsic tissue-scale forces in mice.

**Figure 6.**
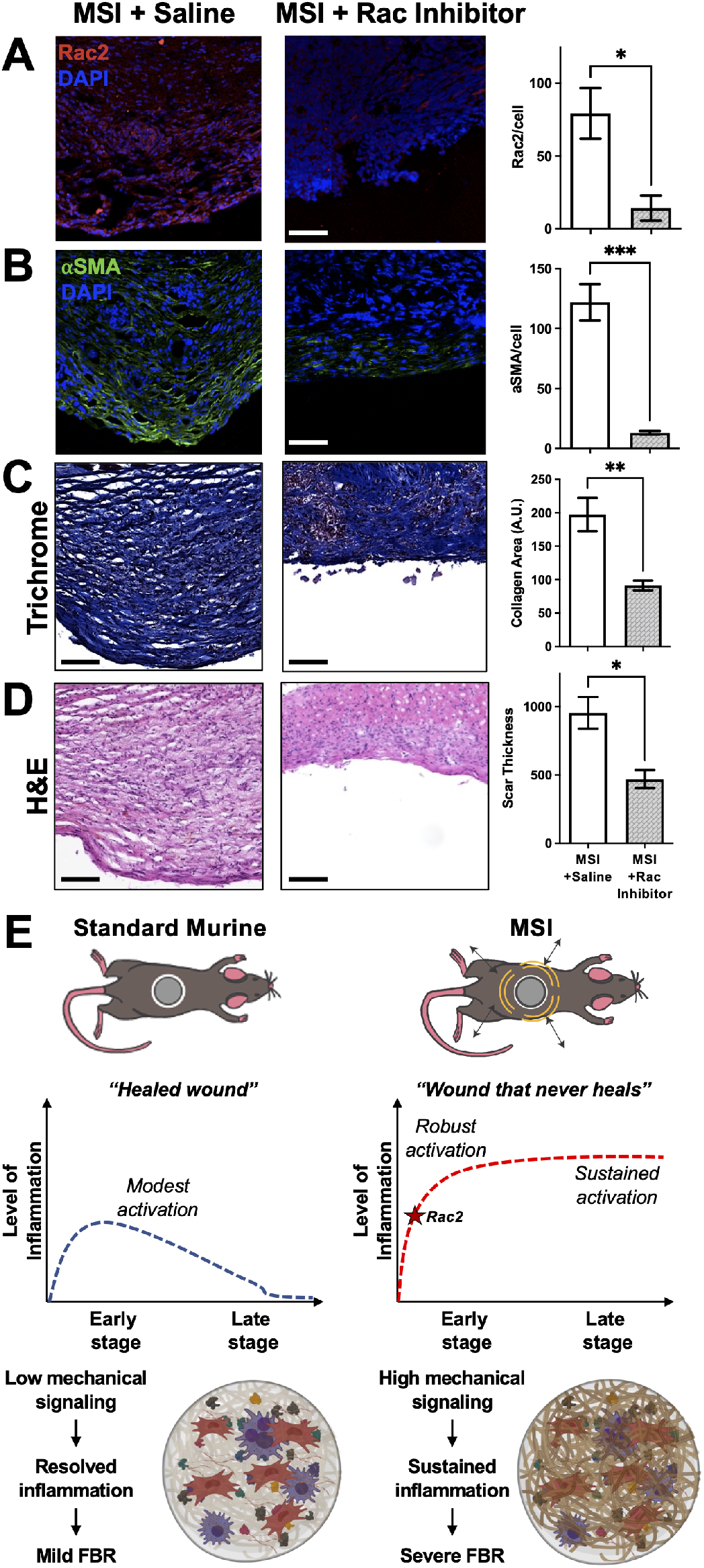
Blocking Rac2 signaling effectively reverses the human-like FBR induced by increased tissue-scale forces in mice. Comparative analysis of histology sections of FBR capsules from the MSI mouse model with and without Rac inhibitor. **(A)** Immunostaining for RAC2 signaling in FBR capsules. Scale bar = 50 μm. Quantification of percent area positive for RAC2 in each capsule. n=5 for each group. *p < 0.05. **(B)** Immunostaining for αSMA signaling in FBR capsules. Scale bar = 50 μm. Quantification of percent area positive for αSMA in each capsule. n=5 for each group. *p < 0.05. **(C)** Trichrome staining of FBR capsules. Scale bar = 50 μm. Quantification of percent area positive for collagen in each capsule. n=5 for each group. *p < 0.05. **(D)** H&E staining of FBR capsules. Scale bar = 50 μm. Quantification of average capsule thickness. n=4 for each group. *p < 0.05. **(E)** In standard murine (SM) implants, there is a modest activation of inflammatory pathways at the early timepoint, which subsides at the late timepoint, resulting in minimal FBR. In contrast, MSI capsules, increased tissue-scale forces lead to the activation of RAC2 mechanical signaling, which promotes a robust activation of inflammatory markers that is sustained over time, resulting in a human-like pathological FBR.

## Discussion

Previous research has identified implant chemistry and mechanical properties as critical factors in mediating FBR (*7, 9*). In this work, we have identified a third, independent variable, extrinsic tissue-scale forces, that play a central role in the foreign body response. In humans, extrinsic tissue-scale forces increase exponentially with body size due to allometric scaling principles, creating a high mechanical stress environment around biomedical implants. In contrast, mice have low extrinsic tissue scale forces, leading to a low mechanical stress environment and minimal FBR. These dramatic changes in tissue forces can explain the stark differences in FBR between murine and humans, independent of both material chemistry and mechanical properties.

Utilizing a novel mechanically stimulating implant, we artificially increased the extrinsic tissue scale forces and recapitulated all aspects of human FBR at both the histological and transcriptional levels. It is important to note that we compared FBR responses to standard murine implants and MSIs using the same material and with exactly the same implant geometry. By varying only tissue-scale forces, we were able to dramatically alter the molecular and cellular response to recreate human-like FBR in mice. Our results provide an explanation for the long-standing conundrum in the field regarding the significant inter-species variability in FBR, despite the fact that the molecular machinery responsible for inflammation and fibrosis is highly conserved among species (*13, 14*). We show that larger-sized organisms, such as humans, experience higher tissue-scale forces because of allometric scaling that then drive subsequent FBR.

The wound healing cascade in humans is characterized by sequential inflammatory and fibrotic phases and eventual quiescence (*67*). FBR begins as a wound healing response to the local tissue damage that occurs during surgical implantation of a device as well, but elevated levels of extrinsic mechanical forces on the implant by the surrounding tissue creates in humans creates a high mechanical stress environment that leads to a sustained inflammatory response. In standard murine implants with 100-fold lower mechanical stress than in humans, we observed a modest inflammatory response at the early stage, which subsided by later timepoints similarly to a “healed wound” (**Fig. 6E**). In contrast, both humans and murine MSIs generate a 100-fold increased stress environment to perpetuate a sustained presence of mechanically activated immune cells at the implant-tissue interface. Thus, increased tissue-scale forces, shaped by allometric scaling principles in humans, result in a “wound that never heals”, which leads to pathological FBR. In fact, allometric tissue-scale forces may provide the missing link explaining other key aspects of pathological FBR, including the activation of fusogenic macrophages, MHC class II lymphocytes, and myofibroblasts.

Importantly, we found that both murine and human pathological FBR capsules were comprised mostly of immune cells and that extrinsic tissue-scale forces result in the mechanical activation of *Rac2* signaling in a unique population of immune cells, with a gene signature conserved in both mice and humans. Most prior literature has focused on mechanoresponsive fibroblasts as the “drivers’ of fibrosis, responding to the highly mechanically stressed environment by secreting cytokines and upregulating inflammatory cell recruitment (*68, 69*). Our findings reveal a more complex interplay between immune cells and fibroblast activity and suggest that mechanoresponsive immune cells under elevated tissue-scale forces may actually drive and regulate the activation of fibroblasts. Some recent reports have indeed emphasized the potentially critical role of immune cells in FBR (*70, 71*). The current findings provide a novel mechanistic link for the activation of immune cells in FBR, namely, *Rac2* signaling in immune cells activated by allometric tissue-scale forces in humans at the implant-tissue interface.

Since allometric tissue properties in humans cannot be altered because they are inherent to the size of the organism, biomolecular and pharmacologic strategies will be required to create truly biointegrative devices. We demonstrated that pharmacological inhibition of *Rac2* could potentially serve as an effective therapy for humans receiving biomedical implants to prevent FBR, increasing patient quality of life, and reducing implant failure rates. Our findings demonstrate a comprehensive characterization to better understand FBR development and identify a mechanistic target to prevent pathological FBR. Collectively, these findings provide novel insights into FBR and have profound implications for the design and safety of all implantable medical devices in humans.

## ACKNOWLEDGEMENTS

We thank Yujin Park for her assistance in tissue processing, Theresa Carlomagno for administrative support, and Dr. Reinhold Dauskardt for helpful discussions. Implant schematics were generated using BioRender.com.

## Author Contributions

Conceptualization: JP, GCG. Methodology: JP, KC, DS, DH, MJ, GCG. Investigation: JP, KC, BK, CB, TD, DH, DS, ZB, PT, HH, JB, NM, MB, AT, MJ, SK, JD, LP, GCG. Visualization: JP, KC, DS, MB, MJ, GCG. Funding acquisition: GCG. Supervision: JP, KC, DS, MJ, GCG. Writing: JP, KC, DS, MJ, JD, LP, GCG.

## Data and Materials Availability

All data to support the manuscript conclusions can be found in the figures and the supplementary materials. Any other data can be requested from the corresponding authors. RNA-seq data will be uploaded to a Github repository and shared during final publication.

## SUPPLEMENTARY MATERIALS

Materials and Methods

Figs. S1 to S10

## MATERIALS AND METHODS

### Human implant capsule specimen

Explanted biomedical devices (breast tissue expanders and implants, neurostimulator batteries, pacemakers, and orthopedic implants) and the surrounding capsular tissue were collected for this study and analyzed. Informed consent was obtained from each patient in accordance with the Institutional Review Board at Stanford University (IRB #41066).

### Human tissue bank and RNA analysis

We employed a large tissue bank for human breast implant capsule tissues, located in Regensburg, Germany, which consists of over 710 unique breast tissue samples (*1*). As relatively few patients with Baker I capsules undergo revisionary surgery, our overall sample size was limited by this group. We were able to identify 9 samples of Baker I capsules in our biobank and this determined the sample size for this study (n=9 for Baker I samples; n = 11 for Baker IV samples). The patients were of comparable ages: i) 40.6±3.9 years at the time of implantation in Baker I and 35.8±4.40 years in Baker IV, and ii) 50.3±3.0 years at the time of explant in Baker I and 51.0±4.0 years in Baker IV. The patients had silicone breast implants placed for augmentation for a mean of 10.7±2.8 years in Baker I and 15.2±4.5 in Baker IV. None of the patients previously had cancer. For RNA analysis, 5μm FFPE sections of human samples were lysed, proteinase K-digested, and analyzed by the HTG EdgeSeq qNPA assay (HTG Molecular Diagnostics, Tucson, AZ) using a biomarker panel (HTG Oncology Biomarker Panel), a 2,549-gene probeset, including markers for inflammation and fibrosis. Following EdgeSeq qNPA processing, samples were individually barcoded by Polymerase Chain Reaction (PCR) and pooled for sequencing. Libraries were sequenced on the Illumina NextSeq platform (Illumina, San Diego, CA) and data was processed with HTG’s parser software. Approval was given by the local ethic committee in Regensburg (Reference No.: 15-101-0024). Differential expression analysis was performed with the EdgeR package in R (v3.14.0) with Benjamini-hochberg correction for multiple hypothesis testing (*2*). The 100 most highly ranked genes from this analysis for Baker I and Baker IV implants were used to perform gene set enrichment analysis against pathway databases using the Database for Annotation, Visualization and Integrated Discovery (DAVID) toolkit as described previously (*3*).

### STRING analysis

To study the interaction of *RAC2* with other Baker IV genes, STRING (Search Tool for the Retrieval of Interacting Genes/Proteins), a pathway analysis tool that predicts genegene interactions was employed as described before (*4*). The minimum required interaction score was set at 0.200. STRING analyzed interactions between the different genes based on experimental evidence as well as the predicted interactions. The relative positions of nodes and the distances between the different nodes are arbitrary. The genegene interactions, which are color-coded as listed below: pink (experimentally determined), blue (curated databases), green (gene neighborhood), red (gene fusion), dark blue (gene co-currence), light green (text mining), black (co-expression), violet (protein homology).

### Mice and Animal Care

All mice used in this study were housed in the Stanford University Veterinary Service Center and NIH and Stanford University animal care guidelines were followed. All procedures were approved by the university’s Administrative Panel on Laboratory Animal Care. C57/BL6 wildtype mice (Jackson labs Stock No: 000664) were used in these experiments.

### Implant fabrication

Standard silicone implants were made of polydimethylsiloxane (PDMS) and fabricated using a Sylgard 184 elastomer base and curing agent as previously described (*5*). A ratio of 5 (Elastomer): 1 (Curing agent) was used for the experiments described. All implants were cylindrical in shape with a 1.55 cm diameter and 0.67+0.07 cm height. For mechanically stimulated implants, a pre-fabricated coin motor (Precision Microdevices) was placed in the elastomer solution before curing (**Fig. 2b**), while controls were PDMS alone. To enable in situ vibration of MSIs, the wires from the implant had to be guided through the skin, which required a novel surgical technique (**Fig. S5**). After skin incision and creation of a subcutaneous pocket on the back of the mice, two 20 G cannulas were inserted into the pocket in a cranio-caudal direction. The wires were tunneled through the pocket and guided through the skin using the cannulas and a modified Seldinger technique, enabling activation of the motor by an external battery. MSIs could then be attached to the external battery for an hour every day during the fibroproliferative phase of FBR (Days 4-11), as outlined in **Fig. S7**. Longer durations of vibration were not well-tolerated by mice. A 3V power source was chosen in accordance with our FE modeling to most accurately match the mechanical stress around implants in humans. As a second control, MSI implants were also tested without *in situ* vibration.

### Implant Mechanical Testing

The Young’s Modulus (E_y_) of silicone implants were determined using a custom compressive test method on Instron 5560 as described previously (*6*). Each sample had a diameter of 1.55cm and subjected to a compressive rate of 1mm/sec. E_y_ of each implant was calculated by taking the linear slope of the stress-strain curve between 0 and 0.10 compressive strain.

### Implantation Experiments

Standard silicone implants and MSIs were implanted in C57/BL6 mice for either 2 weeks or 1 month. A 2 cm incision was made on the dorsum of the mouse and a subcutaneous pocket was created. Control implants were placed in the subcutaneous pocket and the incision was closed using 6-0 nylon suture or staples. For MSIs, the implants were placed into a subcutaneous pocket, similar to the procedure described above, and wires were guided through the skin using a modified seldinger technique. 3V batteries with an amplitude of 1.38G and a frequency of 203Hz were used to mimic human conditions. MSIs were vibrated for 1 hour daily from day 4 (D4) post implantation to day 11 (D11) (**Fig. S7**). This time period was chosen based on previous studies, showing that increased mechanical stress during this period effectively induces fibrosis (*7, 8*).

### Computational Modeling of biomechanical stress patterns around biomedical implants

Computational finite element (FE) models for human and mouse implants were developed using the commercial finite-element software ABAQUS (version 2017, SIMULIA, Providence, RI), using a similar FEM framework as previously described to study mechanical behavior of soft tissues as well as investigate deformation and stress patterns in biological tissues (*9–11*). Model geometry was based on experimental measurements of skin and fat layers and custom designed implants for humans and mice (*12–14*). In the initial configuration, the implants were modeled as a 3D disk (**Fig. S3**). Fat, skin and muscle were represented as layers around the implant. Movement of implants transfers deformation and force to the layers of skin and fat. In all models, the bottom end of the muscle/bone layer was fixed. Tetrahedral elements (C3D4) were used for soft tissue layers, while hexagonal elements (C3D8) were used for implants. Mesh refinement confirmed that the chosen mesh size is accurate enough for the present purposes.

The simulation was based on the theory of elastic deformation of soft tissue where for each human and mouse model, different material properties were considered for skin, fat and muscle/bone layers based on previous findings (**Fig. S3**) (*12, 15, 16*). The models contained external loading as static or vibrating forces, where the direction of applied force was in the horizontal axis in all models. Once the force was applied to implant, this force moves the 3D geometry of the implant in the direction of applied load. Due to the defined tie interaction between implant and tissue layers around it, implant movement applies the stress to the tissue where tissue on both ends of the implant experience negative (compression) and positive (stretch) stresses (**Fig. 2**). The magnitude of these stresses was proportional to the stiffness and elastic properties of the tissue. The stress experienced by the tissue further triggers mechanotransduction pathways, leading to biological responses. The models contained external loading as static or vibrating forces, where the direction of applied force was in the horizontal axis in all models. For human and mouse static models (**Fig. 2A**), the amplitude of the applied force was calculated from dynamic resting tensions reported previously (*7*). For the MSI model (**Fig. 2B**), a periodic force was defined for vibrating implants, where the amplitude of the vibrating force was

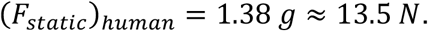

### Histology and Trichrome staining

At each timepoint, the mice were euthanized, and the implants were resected *en bloc* with the surrounding scar tissue. The implants were removed, and the scar tissue along with the skin was fixed in 4% paraformaldehyde overnight and embedded in paraffin. Scar tissue and implants from humans were collected from patients and processed within an hour. For analysis of the capsule, paraffin sections were stained with trichrome (SigmaAldrich) as described previously (*7, 8*). Imaging was performed on a Leica DM5000 B upright microscope. Image analysis software (Image J) was used to quantify collagen staining.

### Herovici’s staining

Herovici’s staining was performed according to manufacturer specifications as described below. Thin histological sections were deparaffinized and immersed in Weigert’s hematoxylin solution for five minutes followed by Herovici’s working solution (equal parts of stain solutions A and B) for 2 minutes. Slides were subsequently immersed in 1% acetic acid, followed by dehydration using alcohol and xylene washes. Finally, slides were mounted using mounting media with a coverslip on top. Imaging was performed on a Leica DM5000 B upright microscope. Image analysis software (Image J) was used to quantify mature collagen (red color) staining.

### Scanning electron microscopy (SEM)

Tissue on the surface of implants was fixed using 4% paraformaldehyde (PFA). Samples were dehydrated using a series of ethanol washes with increasing concentration from 70% to 100% ethanol for five minutes each. The samples were subsequently immersed in hexamethyldisilazane for 15 min and then sputter coated with gold-palladium prior to imaging with SEM. For image analysis, at least eight different SEM images with collagen fibers on the surface of the implant were analyzed using Image J for each group.

### Immunostaining

Immunohistological staining was performed on paraffin sections as described previously (*8, 17*). Briefly, heat-based antigen retrieval was followed by blocking with 5% goat serum in PBS. The following primary antibodies were used at a 1:200 dilution (as recommended by the manufacturer) and incubated overnight at 4°C: anti-αSMA (Abcam ab5694) or anti-Rac2 (Fischer Scientific, DF6273). Incubation of primary antibody-stained specimens with Alexa Fluor 488 secondary antibody (Thermo Fisher Scientific) was performed at a 1:400 dilution for 1 hour at room temperature. Sections were subsequently mounted using Fluoroshield (F6057, Sigma, Saint Louis, MO) with 4’, 6-diamidino-2-phenylindole (DAPI) to stain cell nuclei. Imaging was performed on a Leica DM5000 B upright microscope. Image analysis was performed using MATLAB (*18, 19*).

### Single cell barcoding, library preparation, and single cell RNA sequencing

Implant capsule tissue were obtained from patients undergoing routine implant removal procedures and was processed for single cell sequencing as described previously (*20, 21*). Freshly obtained tissue from the clinic was micro-dissected and digested with collagenase to obtain cellular suspensions for 10x single cell sequencing (Single Cell 3’ v2, 10x Genomics, USA) according to the manufacturer’s instructions. Briefly, a mixture of droplet-based single cell suspensions, partitioning oil, and the reverse transcription master mix were loaded onto a single cell chip, and reverse transcription was performed on the Chromium controller at 53C for 45 mins. cDNA was amplified for 12 cycles on a BioRad C1000 Touch thermocycler using SpriSelect beads (Beckman Coulter, USA) and a ratio of SpriSelect reagent volume to sample volume of 0.6. cDNA was analyzed on an Agilent Bioanalyzer High Sensitivity DNA chip for qualitative control. cDNA was fragmented using the proprietary fragmentation enzyme blend at 32C for 5min, which was followed by end repair and A-tailing for 30min at 65C. cDNA were double-sided size selected using SpriSelect beats, followed by ligation with sequencing adaptors at 20C for 15min. cDNA amplification was performed using a sample-specific index oligo as primer, and subsequently another round of double-sided size selection using SpriSelect beads was performed. Final libraries were analyzed on an Agilent Bioanalyzer High Sensitivity DNA chip for quality control and were sequenced using a HiSeq 500 Illumina platform aiming for 50,000 reads per cell.

### Data processing, FASTQ generation, and read mapping

The Cell Ranger software (10X Genomics; version 3.1) implementation of *mkfastq* was employed to convert base calls to reads. These reads were then aligned against the mm10 v3.0.0 genomes using Cell Ranger’s count function (STAR v2.7.0) with SC2Pv2 chemistry and 5000 expected cells per sample (*22*). Cell barcodes representative of quality cells with at least 200 unique transcripts and less than 10% of their transcriptome of mitochondrial origin were analyzed.

### Data normalization and cell subpopulation identification

Raw Unique molecular identifiers or UMIs from each cell barcode were normalized with a scale factor of 10,000 UMIs per cell. The UMI reads were then natural log transformed with a pseudocount of 1 using the R package Seurat (version 3.1.1) (*23*). Highly variable genes were identified, and cells were scaled by regression to the fraction of mitochondrial transcripts as described previously (*21*). The aggregated data was subsequently evaluated using uniform manifold approximation and projection (UMAP) analysis over the first 15 principal components and cell annotations were ascribed using SingleR toolkit (version 3.11) against the Immgen and mouse RNA-seq databases.

### Generation of characteristic subpopulation markers and enrichment analysis

Seurat’s native *FindMarkers* function with a log fold change threshold of 0.25 using the ROC test was used to generate marker lists for each cluster. The most highly ranked genes from this analysis were used to perform gene set enrichment analysis against pathway databases for each cluster or subgroup of cells using the Database for Annotation, Visualization and Integrated Discovery (DAVID) toolkit as described previously (*3*).

### Rac Inhibition Experiments

EHT 1864 2HCl, a potent Rac family GTPase inhibitor, was acquired from Selleckchem (Houston, TX). MSIs were implanted in C57/BL6 wildtype mice for 28 days utilizing the modified Seldinger technique described above. Mice were injected with EHT 1864 2HCl (10mg/kg/day) (n=4) or saline (n=4) from Days 0-26. FBR capsule tissue was explanted on day 28 and processed for histologic analysis.

### Statistical Analyses

Results are presented as mean ± SEM. Standard data analysis was performed using student’s t-tests. ANOVA with posthoc tukey’s test was used for multiple comparisons. Results were considered significant for *p < 0.05.

## References

1. K. Information, “The Global Market for Medical Devices,” (2019).

2. M. R. Major, V. W. Wong, E. R. Nelson, M. T. Longaker, G. C. Gurtner, The foreign body response: at the interface of surgery and bioengineering. Plast Reconstr Surg 135, 1489–1498 (2015).

3. E. B. Dolan et al., An actuatable soft reservoir modulates host foreign body response. Sci Robot 4, (2019).

4. M. T. Novak, F. Yuan, W. M. Reichert, Modeling the relative impact of capsular tissue effects on implanted glucose sensor time lag and signal attenuation. Anal Bioanal Chem 398, 1695–1705 (2010).

5. S. Mittal et al., Cardiac implantable electronic device infections: incidence, risk factors, and the effect of the AigisRx antibacterial envelope. Heart Rhythm 11, 595–601 (2014).

6. J. M. Anderson, A. Rodriguez, D. T. Chang, Foreign body reaction to biomaterials. Semin Immunol 20, 86–100 (2008).

7. A. J. Vegas et al., Combinatorial hydrogel library enables identification of materials that mitigate the foreign body response in primates. Nat Biotechnol 34, 345–352 (2016).

8. R. A. Bank, Limiting biomaterial fibrosis. Nat Mater 18, 781 (2019).

9. N. Noskovicova et al., Suppression of the fibrotic encapsulation of silicone implants by inhibiting the mechanical activation of pro-fibrotic TGF-beta. Nat Biomed Eng, (2021).

10. L. Zhang et al., Zwitterionic hydrogels implanted in mice resist the foreign-body reaction. Nat Biotechnol 31, 553–556 (2013).

11. V. Yesilyurt et al., A Facile and Versatile Method to Endow Biomaterial Devices with Zwitterionic Surface Coatings. Adv Healthc Mater 6, (2017).

12. J. Padmanabhan et al., In Vivo Models for the Study of Fibrosis. Adv Wound Care (New Rochelle) 8, 645–654 (2019).

13. J. Zhang et al., Molecular Profiling Reveals a Common Metabolic Signature of Tissue Fibrosis. Cell Reports Medicine 1, 100056 (2020).

14. J. Godec et al., Compendium of Immune Signatures Identifies Conserved and Species-Specific Biology in Response to Inflammation. Immunity 44, 194–206 (2016).

15. A. A. Biewener, Scaling body support in mammals: limb posture and muscle mechanics. Science 245, 45–48 (1989).

16. J. R. Hutchinson, M. Garcia, Tyrannosaurus was not a fast runner. Nature 415, 1018–1021 (2002).

17. K. Chen et al., Disrupting biological sensors of force promotes tissue regeneration in large organisms. Nat Commun 12, 5256 (2021).

18. J. C. J. Wei et al., Allometric scaling of skin thickness, elasticity, viscoelasticity to mass for micro-medical device translation: from mice, rats, rabbits, pigs to humans. Scientific Reports 7, 15885 (2017).

19. R. C. Rennert et al., A histological and mechanical analysis of the cardiac lead-tissue interface: implications for lead extraction. Acta Biomater 10, 2200–2208 (2014).

20. B. Kuehlmann, R. Burkhardt, N. Kosaric, L. Prantl, Capsular fibrosis in aesthetic and reconstructive-cancer patients: A retrospective analysis of 319 cases. Clin Hemorheol Microcirc 70, 191–200 (2018).

21. W. Huang da, B. T. Sherman, R. A. Lempicki, Systematic and integrative analysis of large gene lists using DAVID bioinformatics resources. Nat Protoc 4, 44–57 (2009).

22. V. W. Wong, S. Akaishi, M. T. Longaker, G. C. Gurtner, Pushing back: wound mechanotransduction in repair and regeneration. The Journal of investigative dermatology 131, 2186–2196 (2011).

23. S. Joshi et al., Rac2 is required for alternative macrophage activation and bleomycin induced pulmonary fibrosis; a macrophage autonomous phenotype. PLoS One 12, e0182851 (2017).

24. F. Y. McWhorter, C. T. Davis, W. F. Liu, Physical and mechanical regulation of macrophage phenotype and function. Cell Mol Life Sci 72, 1303–1316 (2015).

25. N. J. Meyer et al., GADD45a is a novel candidate gene in inflammatory lung injury via influences on Akt signaling. FASEB J 23, 1325–1337 (2009).

26. S. Mitra et al., Role of growth arrest and DNA damage-inducible alpha in Akt phosphorylation and ubiquitination after mechanical stress-induced vascular injury. Am J Respir Crit Care Med 184, 1030–1040 (2011).

27. K. A. Cho et al., Transplantation of bone marrow cells reduces CCl4-induced liver fibrosis in mice. Liver Int 31, 932–939 (2011).

28. L. Liu et al., Mechanotransduction-modulated fibrotic microniches reveal the contribution of angiogenesis in liver fibrosis. Nat Mater 16, 1252–1261 (2017).

29. K. Chen et al., Mechanical Strain Drives Myeloid Cell Differentiation Toward Proinflammatory Subpopulations. Adv Wound Care (New Rochelle), (2021).

30. Q. Deng, S. K. Yoo, P. J. Cavnar, J. M. Green, A. Huttenlocher, Dual roles for Rac2 in neutrophil motility and active retention in zebrafish hematopoietic tissue. Dev Cell 21, 735–745 (2011).

31. J. L. Dooley et al., Regulation of inflammation by Rac2 in immune complex-mediated acute lung injury. Am J Physiol Lung Cell Mol Physiol 297, L1091–1102 (2009).

32. Y. Cheng, X. L. Ma, Y. Q. Wei, X. W. Wei, Potential roles and targeted therapy of the CXCLs/CXCR2 axis in cancer and inflammatory diseases. Biochim Biophys Acta Rev Cancer 1871, 289–312 (2019).

33. S. Thornton et al., Urokinase plasminogen activator and receptor promote collagen-induced arthritis through expression in hematopoietic cells. Blood Adv 1, 545–556 (2017).

34. Y. Luo et al., Inhibition of macrophage migration inhibitory factor (MIF) as a therapeutic target in bleomycin-induced pulmonary fibrosis rats. Am J Physiol Lung Cell Mol Physiol 321, L6–L16 (2021).

35. D. Szklarczyk et al., STRING v11: protein-protein association networks with increased coverage, supporting functional discovery in genome-wide experimental datasets. Nucleic Acids Res 47, D607–D613 (2019).

36. J. Park et al., A reciprocal regulatory circuit between CD44 and FGFR2 via c-myc controls gastric cancer cell growth. Oncotarget 7, 28670–28683 (2016).

37. S. E. Byeon et al., The role of Src kinase in macrophage-mediated inflammatory responses. Mediators Inflamm 2012, 512926 (2012).

38. L. F. Ferrari, D. Araldi, O. Bogen, J. D. Levine, Extracellular matrix hyaluronan signals via its CD44 receptor in the increased responsiveness to mechanical stimulation. Neuroscience 324, 390–398 (2016).

39. W. H. Guo, M. T. Frey, N. A. Burnham, Y. L. Wang, Substrate rigidity regulates the formation and maintenance of tissues. Biophys J 90, 2213–2220 (2006).

40. K. Chen et al., Role of boundary conditions in determining cell alignment in response to stretch. Proceedings of the National Academy of Sciences of the United States of America 115, 986–991 (2018).

41. K. L. Helton, B. D. Ratner, N. A. Wisniewski, Biomechanics of the sensor-tissue interface-effects of motion, pressure, and design on sensor performance and the foreign body response-part I: theoretical framework. J Diabetes Sci Technol 5, 632–646 (2011).

42. V. W. Wong et al., Focal adhesion kinase links mechanical force to skin fibrosis via inflammatory signaling. Nat Med 18, 148–152 (2011).

43. K. Ren, R. Torres, Role of interleukin-1beta during pain and inflammation. Brain Res Rev 60, 57–64 (2009).

44. L. Backdahl, M. Aoun, U. Norin, R. Holmdahl, Identification of Clec4b as a novel regulator of bystander activation of auto-reactive T cells and autoimmune disease. PLoS Genet 16, e1008788 (2020).

45. N. Choudhry et al., The complement factor 5a receptor 1 has a pathogenic role in chronic inflammation and renal fibrosis in a murine model of chronic pyelonephritis. Kidney Int 90, 540–554 (2016).

46. D. Aran et al., Reference-based analysis of lung single-cell sequencing reveals a transitional profibrotic macrophage. Nat Immunol 20, 163–172 (2019).

47. E. O. Apostolov, X. Wang, S. V. Shah, A. G. Basnakian, Role of EndoG in development and cell injury. Cell Death Differ 14, 1971–1974 (2007).

48. O. Lytovchenko, E. R. S. Kunji, Expression and putative role of mitochondrial transport proteins in cancer. Biochim Biophys Acta Bioenerg 1858, 641–654 (2017).

49. C. Rouault et al., Roles of chemokine ligand-2 (CXCL2) and neutrophils in influencing endothelial cell function and inflammation of human adipose tissue. Endocrinology 154, 1069–1079 (2013).

50. T. T. Chang, J. W. Chen, Emerging role of chemokine CC motif ligand 4 related mechanisms in diabetes mellitus and cardiovascular disease: friends or foes? Cardiovasc Diabetol 15, 117 (2016).

51. P. J. Brooks, M. Glogauer, C. A. McCulloch, An Overview of the Derivation and Function of Multinucleated Giant Cells and Their Role in Pathologic Processes. Am J Pathol 189, 1145–1158 (2019).

52. P. Gonzalo et al., MT1-MMP is required for myeloid cell fusion via regulation of Rac1 signaling. Dev Cell 18, 77–89 (2010).

53. O. Petillo et al., In vivo induction of macrophage Ia antigen (MHC class II) expression by biomedical polymers in the cage implant system. J Biomed Mater Res 28, 635–646 (1994).

54. D. Voehringer, T. A. Reese, X. Huang, K. Shinkai, R. M. Locksley, Type 2 immunity is controlled by IL-4/IL-13 expression in hematopoietic non-eosinophil cells of the innate immune system. J Exp Med 203, 1435–1446 (2006).

55. S. Mascharak et al., Preventing Engrailed-1 activation in fibroblasts yields wound regeneration without scarring. Science 372, (2021).

56. A. Birnhuber, V. Biasin, D. Schnoegl, L. M. Marsh, G. Kwapiszewska, Transcription factor Fra-2 and its emerging role in matrix deposition, proliferation and inflammation in chronic lung diseases. Cell Signal 64, 109408 (2019).

57. R. Rabieian et al., Plasminogen Activator Inhibitor Type-1 as a Regulator of Fibrosis. J Cell Biochem 119, 17–27 (2018).

58. H. Lazareth et al., The tetraspanin CD9 controls migration and proliferation of parietal epithelial cells and glomerular disease progression. Nat Commun 10, 3303 (2019).

59. E. Vidak, U. Javorsek, M. Vizovisek, B. Turk, Cysteine Cathepsins and their Extracellular Roles: Shaping the Microenvironment. Cells 8, (2019).

60. S. P. Turunen, O. Tatti-Bugaeva, K. Lehti, Membrane–type matrix metalloproteases as diverse effectors of cancer progression. Biochim Biophys Acta Mol Cell Res 1864, 1974–1988 (2017).

61. R. Li et al., Pdgfra marks a cellular lineage with distinct contributions to myofibroblasts in lung maturation and injury response. Elife 7, (2018).

62. B. A. Shook et al., Myofibroblast proliferation and heterogeneity are supported by macrophages during skin repair. Science 362, (2018).

63. F. C. Yang et al., Rac and Cdc42 GTPases control hematopoietic stem cell shape, adhesion, migration, and mobilization. Proc Natl Acad Sci U S A 98, 5614–5618 (2001).

64. I. S. Kang, J. S. Jang, C. Kim, Opposing roles of hematopoietic-specific small GTPase Rac2 and the guanine nucleotide exchange factor Vav1 in osteoclast differentiation. Sci Rep 10, 7024 (2020).

65. A. Troeger, D. A. Williams, Hematopoietic–specific Rho GTPases Rac2 and RhoH and human blood disorders. Exp Cell Res 319, 2375–2383 (2013).

66. A. Shutes et al., Specificity and mechanism of action of EHT 1864, a novel small molecule inhibitor of Rac family small GTPases. J Biol Chem 282, 35666–35678 (2007).

67. G. C. Gurtner, S. Werner, Y. Barrandon, M. T. Longaker, Wound repair and regeneration. Nature 453, 314–321 (2008).

68. N. C. Henderson, F. Rieder, T. A. Wynn, Fibrosis: from mechanisms to medicines. Nature 587, 555–566 (2020).

69. R. R. Driskell et al., Distinct fibroblast lineages determine dermal architecture in skin development and repair. Nature 504, 277–281 (2013).

70. L. Chung et al., Interleukin 17 and senescent cells regulate the foreign body response to synthetic material implants in mice and humans. Sci Transl Med 12, (2020).

71. J. C. Doloff et al., Colony stimulating factor-1 receptor is a central component of the foreign body response to biomaterial implants in rodents and non-human primates. Nat Mater 16, 671–680 (2017).

## Supplementary References

1. B. Kuehlmann, R. Burkhardt, N. Kosaric, L. Prantl, Capsular fibrosis in aesthetic and reconstructive-cancer patients: A retrospective analysis of 319 cases. Clin Hemorheol Microcirc 70, 191–200 (2018).

2. M. D. Robinson, D. J. McCarthy, G. K. Smyth, edgeR: a Bioconductor package for differential expression analysis of digital gene expression data. Bioinformatics 26, 139–140 (2010).

3. W. Huang da, B. T. Sherman, R. A. Lempicki, Systematic and integrative analysis of large gene lists using DAVID bioinformatics resources. Nat Protoc 4, 44–57 (2009).

4. D. Szklarczyk et al., STRING v11: protein-protein association networks with increased coverage, supporting functional discovery in genome-wide experimental datasets. Nucleic Acids Res 47, D607–D613 (2019).

5. R. N. Palchesko, L. Zhang, Y. Sun, A. W. Feinberg, Development of polydimethylsiloxane substrates with tunable elastic modulus to study cell mechanobiology in muscle and nerve. PLoS One 7, e51499 (2012).

6. K. Ma et al., Controlled Delivery of a Focal Adhesion Kinase Inhibitor Results in Accelerated Wound Closure with Decreased Scar Formation. J Invest Dermatol 138, 2452–2460 (2018).

7. S. Aarabi et al., Mechanical load initiates hypertrophic scar formation through decreased cellular apoptosis. FASEB J 21, 3250–3261 (2007).

8. V. W. Wong et al., Focal adhesion kinase links mechanical force to skin fibrosis via inflammatory signaling. Nat Med 18, 148–152 (2011).

9. H. S. Hosseini, K. E. Garcia, L. A. Taber, A new hypothesis for foregut and heart tube formation based on differential growth and actomyosin contraction. Development 144, 2381–2391 (2017).

10. H. S. Hosseini, D. C. Beebe, L. A. Taber, Mechanical effects of the surface ectoderm on optic vesicle morphogenesis in the chick embryo. J Biomech 47, 3837–3846 (2014).

11. H. S. Hosseini, L. A. Taber, How mechanical forces shape the developing eye. Prog Biophys Mol Biol 137, 25–36 (2018).

12. J. C. J. Wei et al., Allometric scaling of skin thickness, elasticity, viscoelasticity to mass for micro-medical device translation: from mice, rats, rabbits, pigs to humans. Sci Rep 7, 15885 (2017).

13. K. Calabro, A. Curtis, J. R. Galarneau, T. Krucker, I. J. Bigio, Gender variations in the optical properties of skin in murine animal models. J Biomed Opt 16, 011008 (2011).

14. M. Geerligs, Skin layer mechanics. (2009).

15. I. Jalilian et al., Cell elasticity is regulated by the tropomyosin isoform composition of the actin cytoskeleton. PLoS One 10, e0126214 (2015).

16. N. Shoham, A. Gefen, The Biomechanics of Fat: From Tissue to a Cell Scale. (2016), pp. 79–92.

17. Y. Rinkevich et al., Skin fibrosis. Identification and isolation of a dermal lineage with intrinsic fibrogenic potential. Science 348, aaa2151 (2015).

18. K. Chen et al., Role of boundary conditions in determining cell alignment in response to stretch. Proceedings of the National Academy of Sciences of the United States of America 115, 986–991 (2018).

19. K. Chen et al., Disrupting biological sensors of force promotes tissue regeneration in large organisms. Nat Commun 12, 5256 (2021).

20. M. Januszyk et al., Characterization of Diabetic and Non-Diabetic Foot Ulcers Using Single-Cell RNA-Sequencing. Micromachines (Basel) 11, (2020).

21. T. Leavitt et al., Prrx1 Fibroblasts Represent a Pro-fibrotic Lineage in the Mouse Ventral Dermis. Cell Rep 33, 108356 (2020).

22. A. Dobin et al., STAR: ultrafast universal RNA-seq aligner. Bioinformatics 29, 15–21 (2013).

23. T. Stuart et al., Comprehensive Integration of Single-Cell Data. Cell 177, 1888–1902 e1821 (2019).

