## Supplementary Figures for "Allometric tissue-scale forces activate mechanoresponsive immune cells to drive pathological foreign body response to biomedical implants"

### Supplementary Figure S1

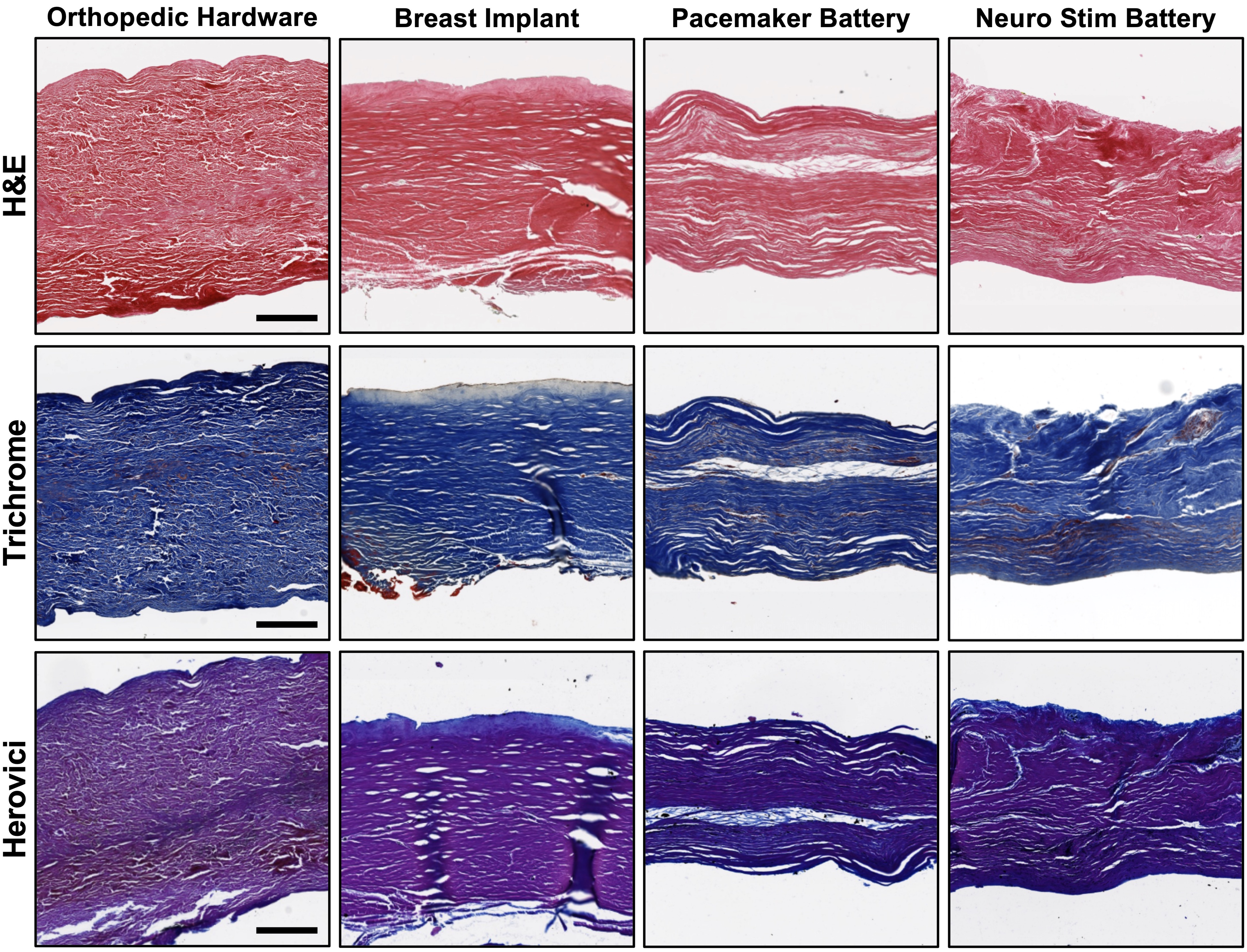


**Figure S1. Pathological, severe FBR in humans is characterized by similar fibrotic encapsulation, regardless of implant properties. (A)** Hematoxylin and Eosin, **(B)** Trichrome, and **(C)** Herovici staining of fibrotic capsules from the fibrous capsule formed around silicone-based breast implants, titanium-based pacemakers, and stainless steel-based orthopedic implants are all strikingly similar to each other on the tissue architectural level.

### Supplementary Figure S2

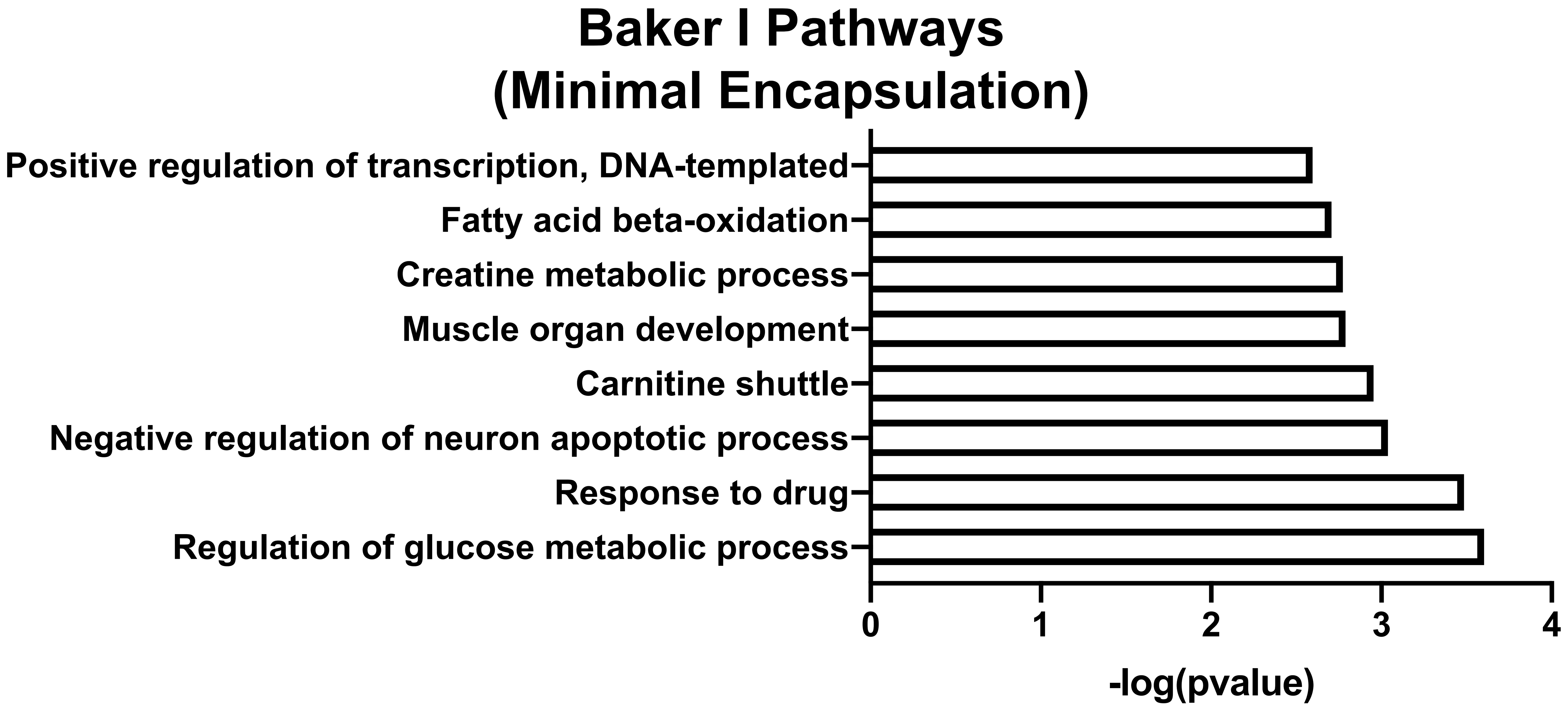


**Figure S2. Pathways significantly upregulated in Baker I samples analyzed using Database for Annotation, Visualization and Integrated Discovery (DAVID).**

### Supplementary Figure S3

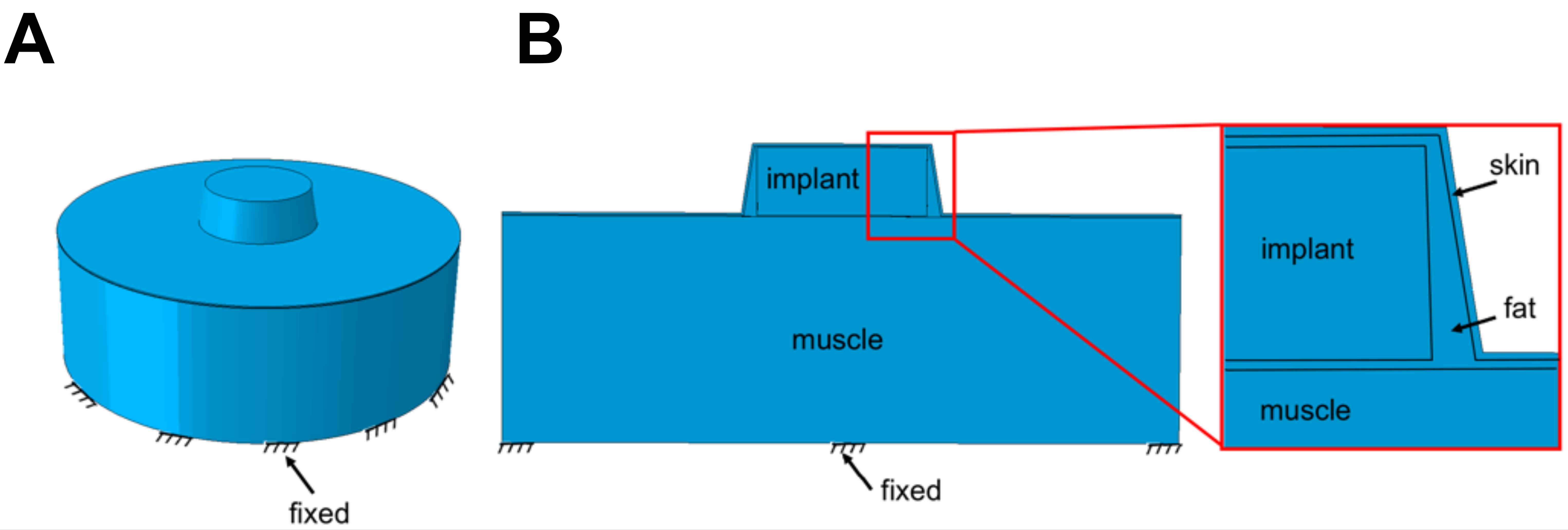


**C** **Dimensions of human and mouse tissue and implants**

| | Human Model ($\mu m$) | Mouse Model ($\mu m$) |
|---|---|---|
| **Thickness of skin** | 2400 | 200 |
| **Thickness of fat-top** | 500 | 100 |
| **Thickness of fat-bottom** | 500 | 100 |
| **Thickness of fat-upper side** | 500 | 100 |
| **Thickness of fat-lower side** | 1000 | 200 |

| | Human Model ($cm$) | Mouse Model ($cm$) |
|---|---|---|
| **Height of Pacemaker** | 0.75 | NA |
| **Radius of Pacemaker** | 2.5 | NA |
| **Height of Chest battery** | 1.5 | NA |
| **Radius of Chest battery** | 2.5 | NA |
| **Radius of Breast Implant** | 12 | NA |
| **Height of Implant** | NA | 0.67 |
| **Radius of Implant** | NA | 0.75 |

**D** **Material properties (Young's modulus) of human and mouse models**

| | Human Model | Mouse Model |
|---|---|---|
| **Skin** | 108.19 KPa | 13.22 KPa |
| **Fat** | 3.25 KPa | 3.5 KPa |
| **Muscle** | 18.6 GPa | 8.9 GPa |
| **Control Implant** | NA | 2.75 MPa |
| **MSI** | NA | 7.80 MPa |
| **Pacemaker** | 103 GPa | NA |
| **Chest battery** | 103 GPa | NA |
| **Breast Implant** | 12 MPa | NA |

**Figure S3. Geometrical model and parameters used for FE modeling of mechanical stress around biomedical implants. (A,B)** Geometrical model used for FE modeling of biomedical implants. **(C)** Average values for skin and subcutaneous tissue properties used for modeling based on previous literature (Refs 61-63).

#### Supplementary Figure S4

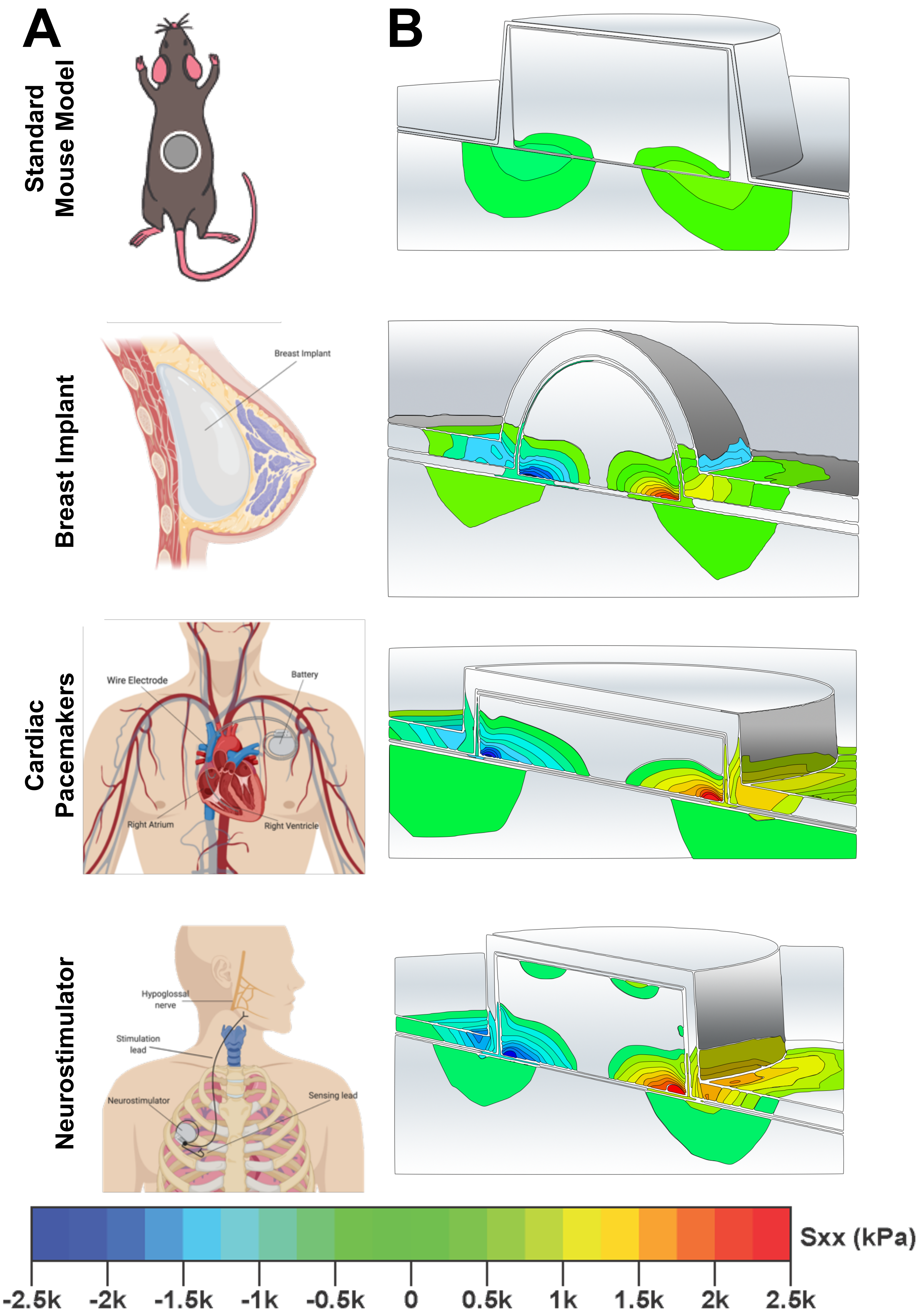


**Figure S4. All human implants experience about 100-fold increased mechanical stress compared to standard murine implants. (A)** Schematic of standard murine implants and and three commonly used human implants: breast implants, pacemakers, and neurostimulator batteries. **(B)** FE modeling of standard murine models of FBR reveals minimal mechanical stress at the implant-tissue interface. FE modeling reveals high mechanical stress (~100-fold higher) around human implants including breast implants, pacemakers and neurostimulator batteries.

### Supplementary Figure S5

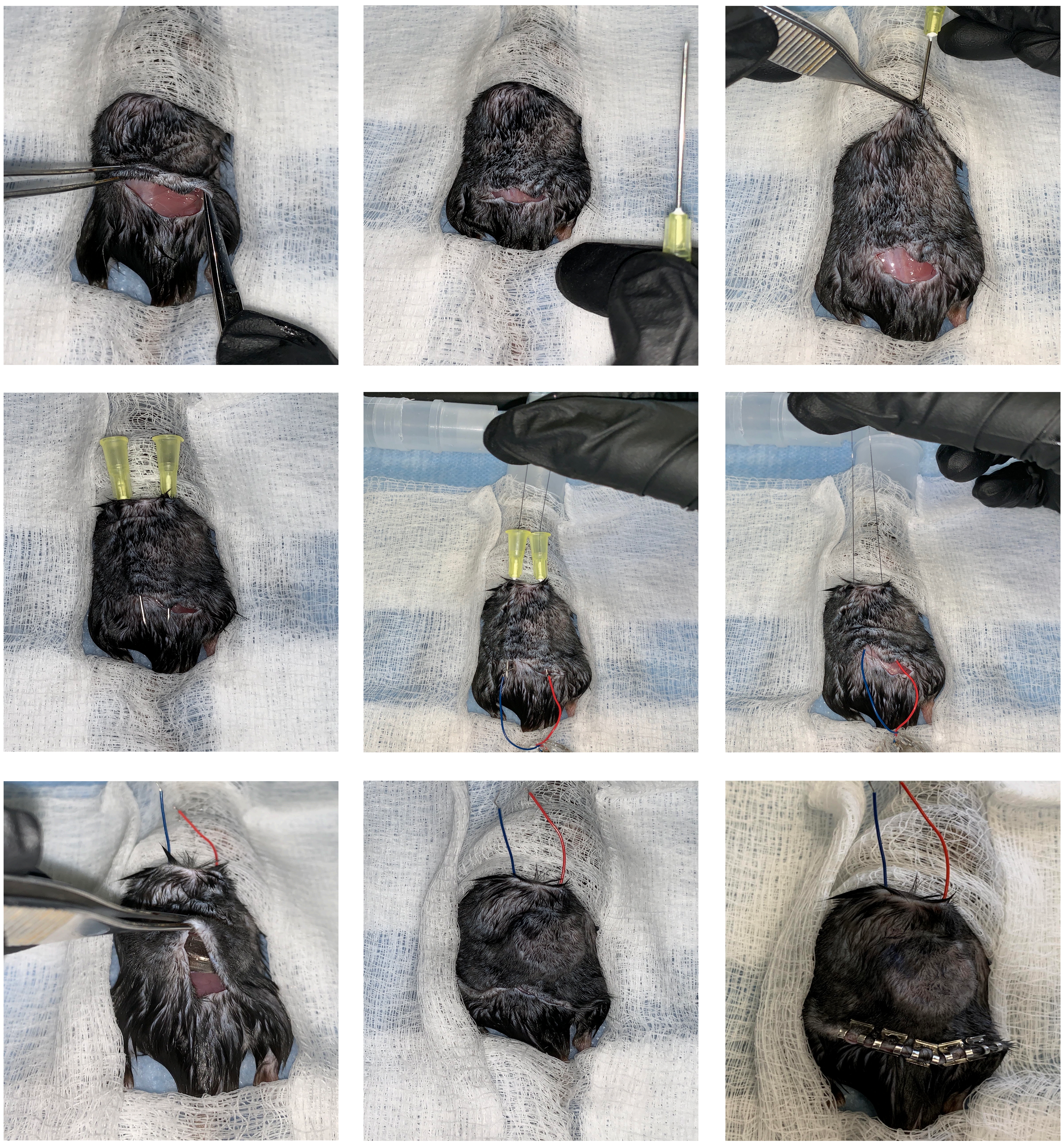

**Figure S5. Method to implant mechanically stimulated implants (MSI).** To induce human levels of mechanical stress in a mouse, we developed silicone implants with an encapsulated pre-fabricated coin motor, capable of in situ vibration. To enable in situ vibration of MSIs, the wires from the implant had to be guided through the skin, which required a novel surgical technique. After skin incision and creation of a subcutaneous pocket on the back of the mice, two 20 G cannulas were inserted into the pocket in a cranio-caudal direction. The wires were tunneled through the pocket and guided through the skin using the cannulas and a modified Seldinger technique, enabling activation of the motor by an external battery.

### Supplementary Figure S6

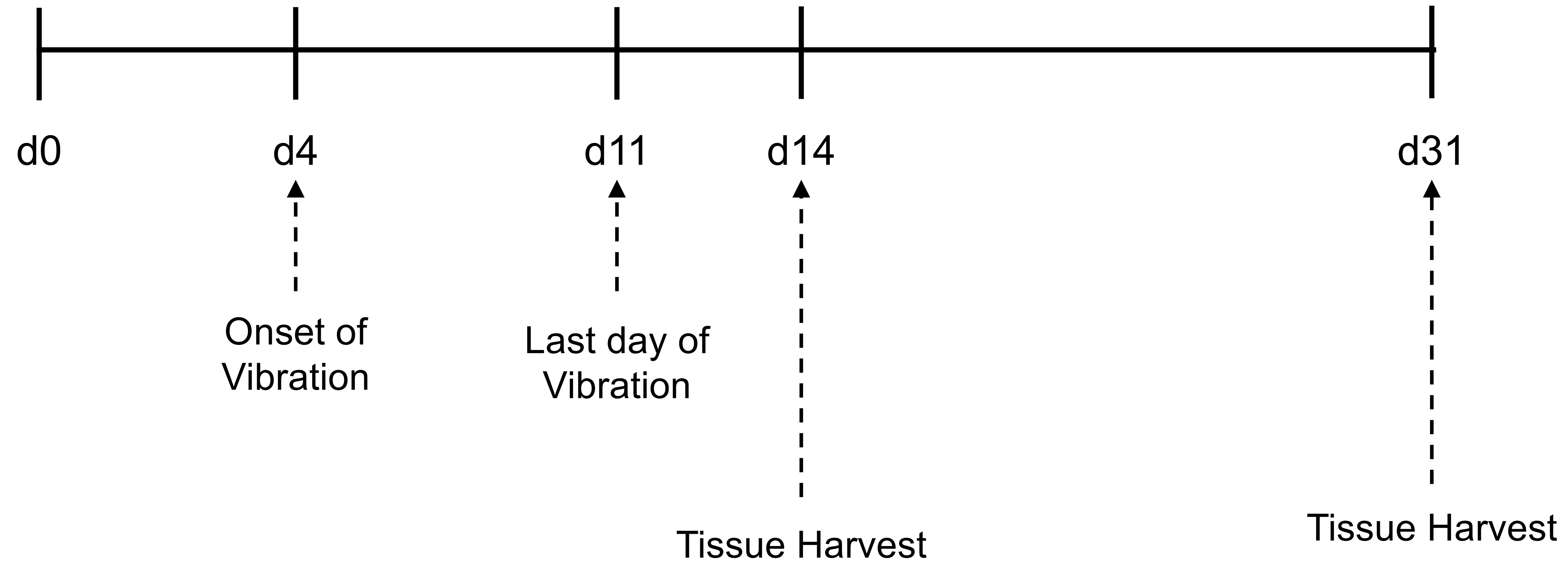


**Figure S6. Timeline of vibration for MSIs.** MSIs were vibrated daily for 1 hour from day 4 to day 12. After iterating through combinations of vibration frequency and amplitude in our finite element model, we determined that 3V batteries with an amplitude of 1.38G and a frequency of 203 Hz would artificially increase extrinsic forces from the surrounding host tissue to generate a 100-fold increase in mechanical stress at the implant-tissue interface (24.1 kPa), similar to that surrounding human implants

### Supplementary Figure S7

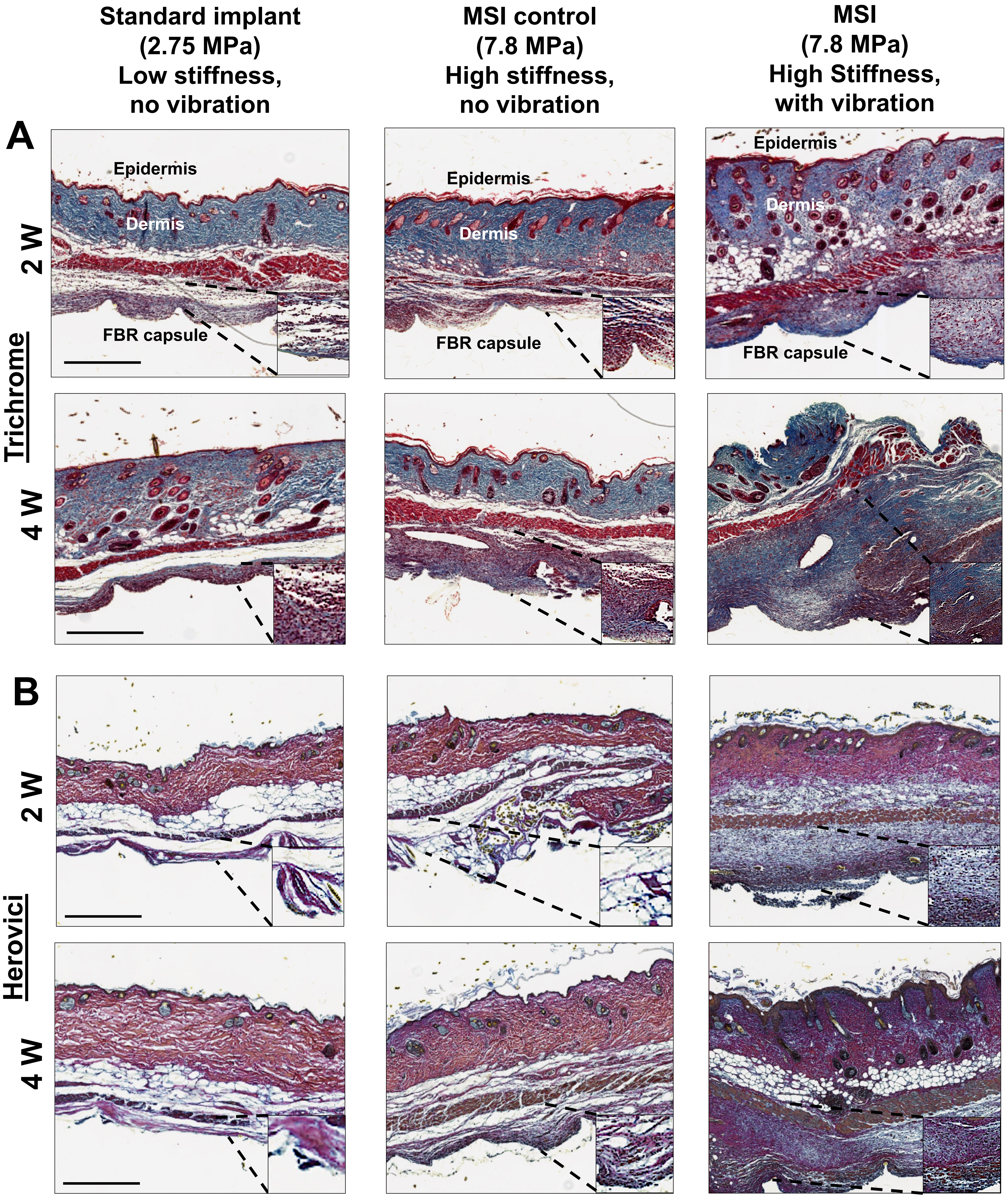


**Figure S7. Increased tissue-scale forces result in increased fibrosis around implants in mice, independent of implant chemistry or mechanical properties. (A,B)** Trichrome staining and Herovici staining of FBR capsules formed around standard murine implants with low stiffness, standard implants with high stiffness, and MSIs reveals that MSI-model produces a more robust scar tissue, with increased collagen and mature collagen. Scale bar = 500μm.

### Supplementary Figure S8

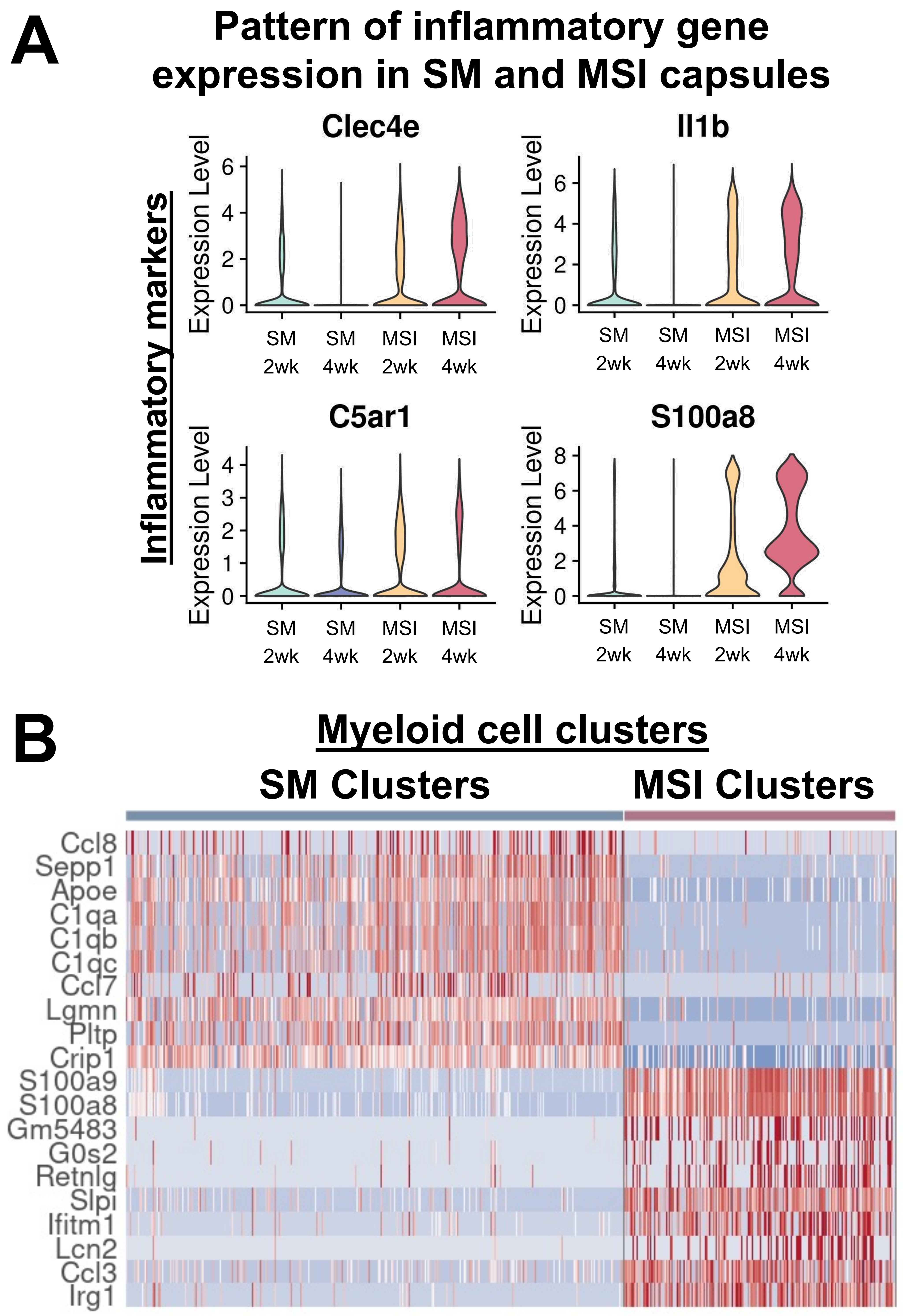


**Figure S8. scRNA-seq of cells from SM and MSI implant capsules. (A)** In standard murine (SM) implants, there is a modest activation of inflammatory pathways at the early timepoint, which subsides at the late timepoint. In contrast, MSI capsules show a robust activation of inflammatory markers that is sustained over time. **(B)** Heatmap of differentially regulated genes in myeloid cells. MSI myeloid cells upregulated inflammatory markers.

### Supplementary Figure S9

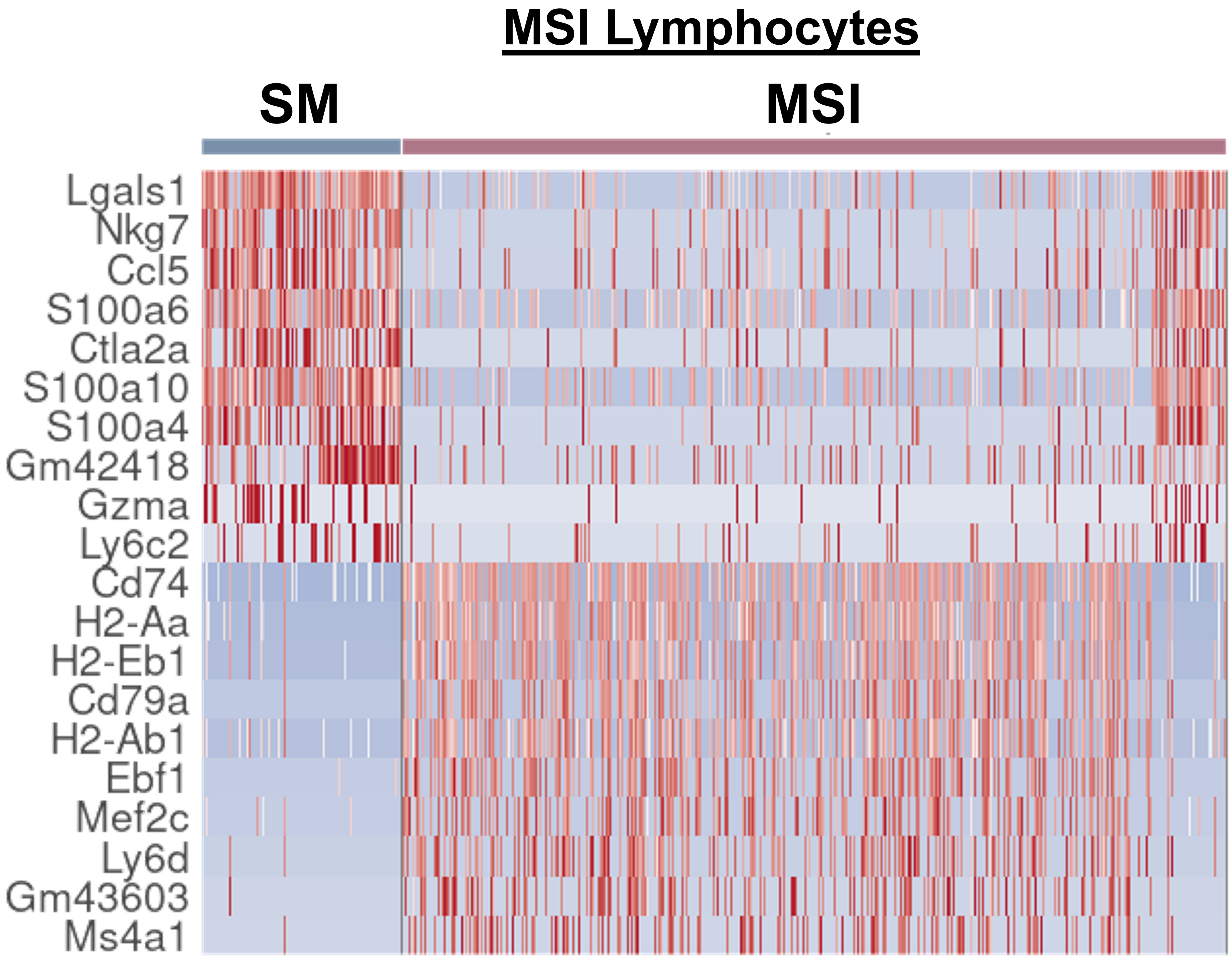


**Figure S9.** Differentially upregulated genes between lymphocytes cells from the standard murine implants and MSIs.

### Supplementary Figure S10

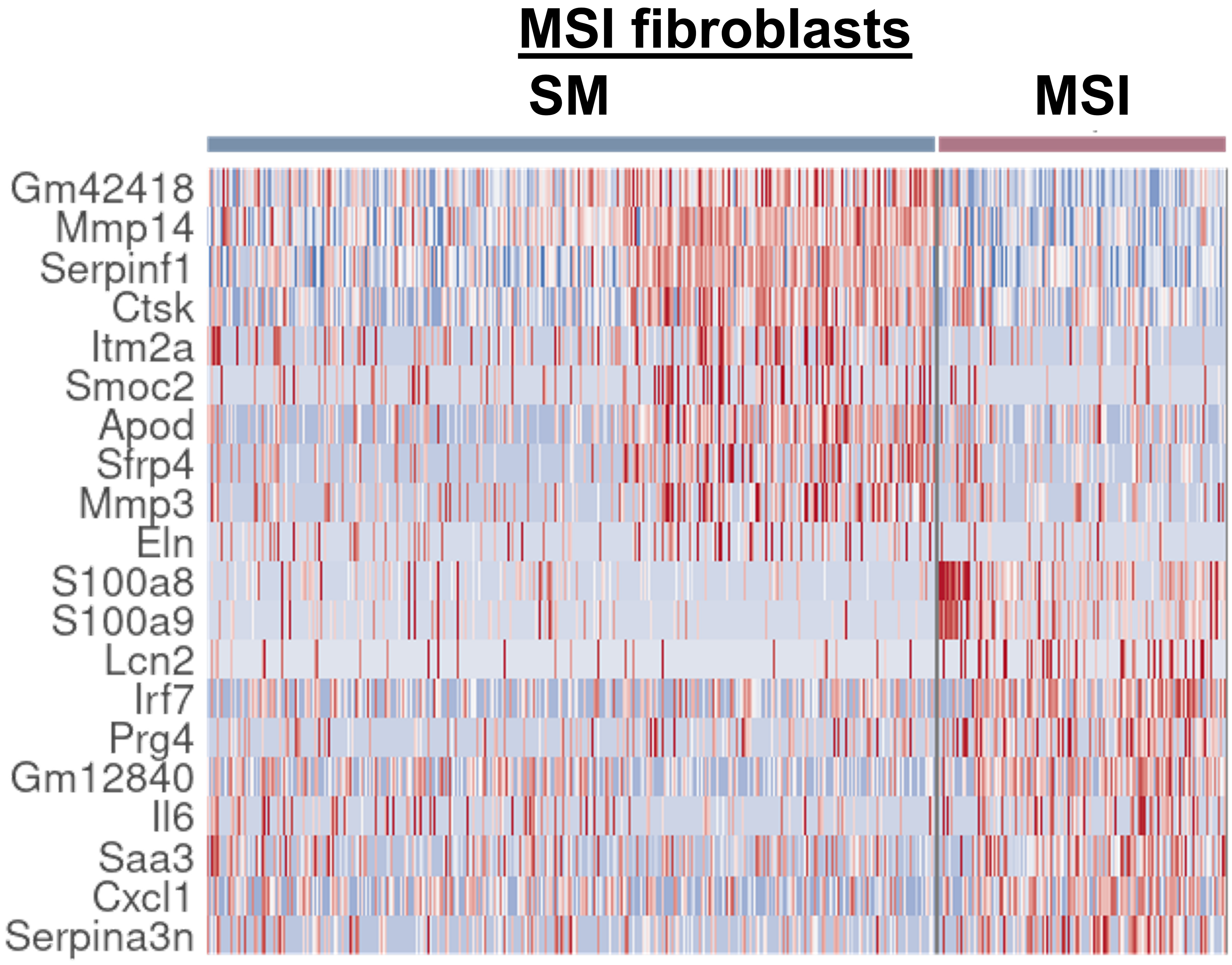


**Figure S10.** Differentially upregulated genes between fibroblast cells from the standard murine implants and MSIs.
